# MIRAGE: a Bayesian statistical method for gene-level rare variant analysis incorporating functional annotations

**DOI:** 10.1101/828061

**Authors:** Shengtong Han, Xiaotong Sun, Laura Sloofman, F. Kyle Satterstrom, Xizhi Xu, Lifan Liang, Nicholas Knoblauch, Wenhui Sheng, Siming Zhao, Tan-Hoang Nguyen, Gao Wang, Autism Sequencing Consortium, Joseph Buxbaum, Xin He

**Affiliations:** School of Dentistry, Marquette University, Milwaukee, WI, USA; Department of Human Genetics, University of Chicago, Chicago, IL, USA; Seaver Autism Center for Research and Treatment, Icahn School of Medicine at Mount Sinai, New York, NY, USA; Department of Psychiatry, Icahn School of Medicine at Mount Sinai, New York, NY, USA; Program in Medical and Population Genetics, Broad Institute of MIT and Harvard, Cambridge, MA, USA; Stanley Center for Psychiatric Research, Broad Institute of MIT and Harvard, Cambridge, MA, USA; Analytic and Translational Genetics Unit, Department of Medicine, Massachusetts General Hospital, Boston, MA, USA; Department of Mathematical and Statistical Sciences, Marquette University, Milwaukee, WI, USA; Department of Biomedical Data Science, Dartmouth College, Hanover, NH, USA; Virginia Institute for Psychiatric and Behavioral Genetics, Virginia Commonwealth University, Richmond, VA, USA; Department of Neurology, Columbia University Vagelos College of Physicians and Surgeons, New York City, NY, USA; Grossman Institute for Neuroscience, Quantitative Biology and Human Behavior, University of Chicago, Chicago, IL, USA

## Abstract

Rare variant analysis is commonly used in whole-exome or genome sequencing studies. Compared to common variants, rare variants tend to have larger effect sizes and often directly point out causal genes. These potential benefits make association analysis with rare variants a priority for human genetics researchers. To improve the power of such studies, numerous methods have been developed to aggregate information of all variants of a gene. However, these gene-based methods often make unrealistic assumptions, e.g. the commonly used burden test effectively assumes that all variants chosen in the analysis have the same effects. In practice, current methods are often underpowered. We propose a Bayesian method: MIxture model based Rare variant Analysis on GEnes (MIRAGE). MIRAGE analyzes summary statistics, i.e. variant counts from inherited variants in trio-sequencing or from ancestry-matched case-control studies. MIRAGE captures the heterogeneity of variant effects by treating all variants of a gene as a mixture of risk and non-risk variants, and uses external information of variants to model the prior probabilities of being risk variants. We demonstrate in both simulations and analysis of an exome-sequencing dataset of autism, that MIRAGE significantly outperforms current methods for rare variant analysis. The top genes identified by MIRAGE are highly enriched with known or plausible autism risk genes. MIRAGE is available at https://xinhe-lab.github.io/mirage.

## 1 Introduction

Genome-wide association studies (GWAS) have identified many loci associated with various complex traits^1–3^. However, identifying causal variants and their target genes is often difficult. Additionally, most common variants discovered by GWAS have small effect sizes^2,3^. These limitations make it difficult to translate GWAS findings into molecular mechanisms of diseases. Sequencing studies focusing on rare variants have the potential to address these challenges. Large effect variants, because of purifying selection, are often rare in the population^4–7^. Indeed, rare variants were estimated to explain between 24% to 50% of heritability of complex traits^8,9^. Another benefit of rare variant analysis is that linkage disequilibrium (LD) is much weaker^10^, making it straightforward to identify causal variants. Furthermore, reduced sequencing cost has made it feasible to sequence large numbers of exomes or genomes. Thus, discovering rare variants underlying the risks of complex diseases from sequencing studies is an important area in human genetics^11^.

Existing rare variant studies often focus on the protein-coding portions of the genome, using Whole Exome Sequencing (WES). WES studies have achieved notable successes in medically important traits such as Alzheimer’s disease and schizophrenia^12–14^. Nevertheless, these studies typically discovered relatively few variants associated with the traits. A natural strategy to improve the power of rare variant studies is to aggregate information from all rare variants in a gene to test if they collectively associate with the phenotype^15^. Many methods have been developed to perform variant set tests^11^. The most commonly used method is the burden test, which collapses all rare, potentially deleterious variants in a gene, and tests the association of the variant “burden” with a phenotype^16–20^. The Sequence Kernel Association Test (SKAT)^21^ assesses if the variance of the effect sizes of the variants is equal to 0. Other tests combine the p values of individual variants, through Fisher’s or similar methods^19,22^, Aggregated Cauchy association test (ACAT)^23^, or the Generalized Berk-Jones (GBJ) test^24^. Additionally, rare variant analysis methods have been developed to incorporate functional information of variants and to deal with unbalanced case-control data^25,26^.

Despite these efforts, the rare variant set test have limited successes. In a WES study of 450,000 subjects from UK Biobank, the burden test found few risk genes for many common traits, for example, 1 gene for Body Mass Index, 3 genes for Type 2 Diabetes and 2 genes for asthma^27^. Often, the power of these tests is lower than the single-variant association test^28^. In a Crohn’s disease WES, researchers found 45 rare risk variants from single-variant analysis, but only 2 risk genes from the burden analysis. These results suggest that causal variants are sparse even within risk genes, leading to a loss of power when collapsing all rare variants in a gene together. Although p-value combination methods such as ACAT were designed to be more sensitive when a small number of variants have causal effects, they do not explicitly model the heterogeneity of effects across variants and continue to collapse variants with frequencies below a threshold^23^. Researchers have attempted to improve the power of rare variant analysis by predicting likely deleterious variants and then running association analysis on these variants, but bioinformatic predictions remain imperfect.

We aim to better model variant effects with a mixture method. Our statistical model can be applied to any settings where we have variant counts from cases, and from population-matched control samples. In other words, it is designed for situations which have properly controlled for population stratification. This simplifies the statistical model and allows us to focus on the challenge of modeling variant effects. In practice, one can obtain such datasets through population-matched controls^29,30^ or through family-based trio studies. In the former, researchers project the genetic data from both cases and potential control subjects into the Principal Component (PC) space, and then select controls that best match the cases in this space. In the latter, variants from parents can be classified as transmitted to children or not. For any variant not associated with the case phenotype, the ratio of transmitted vs un-transmitted alleles should converge to one as sample sizes grow large. Following transmission disequilibrium test (TDT)^31,32^, an imbalanced ratio of a variant would suggest association of the variant with the trait^33^. Thus one can treat transmitted variants as the variant count from cases, and the un-transmitted ones as controls (pseudo-control). Indeed, trio-sequencing studies of autism spectrum disorder (ASD) has revealed an overall tendency of transmission of deleterious variants^34^, but few individual genes have been identified using inherited data, highlighting the need of better data analysis methods.

We propose a Bayesian statistical method to better account for the heterogeneity of the variant effects in a gene or a genomic region. We model the variants in a gene as a mixture of risk and non-risk variants. The prior probability of a variant being a risk variant depends on the functional annotations of the variant. These prior probabilities are generally low, reflecting the sparsity of risk variants, but also vary considerably across variants based on their likely functional effects. This Bayesian strategy of incorporating functional information as prior has significant advantages over simply filtering variants based on their likely effects. In general, the external annotations have limited accuracy in predicting functional effects; and even if a variant is functional, it does not necessarily have an effect on the particular trait of interest. Importantly, the parameters linking the annotations of variants and their prior probabilities are estimated using an Empirical Bayes strategy, by pooling data from all genes. Our model has the additional advantage that it requires only summary statistics (variant counts). This makes it computationally efficient and eliminates the need of sharing individual-level data.

We demonstrated the advantage of the proposed method, MIxture model based Rare variant Analysis on GEnes (MIRAGE), in detecting putative risk genes over existing methods, in both simulation studies, and the analysis of inherited variants from ASD whole exome sequencing studies.

## 2 Methods

### 2.1 MIRAGE model

The input data of MIRAGE include rare variant counts in cases and controls with sample sizes *N*_1_ and *N*_0_, respectively. For variant *j* of gene *i*, we denote *X*_*ij*_ its allele count in cases, 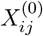 in controls, and *T*_*ij*_ its total allele counts in cases and controls. We also have annotations of each variant. We assume each variant belongs to one variant category, denoted as *c*_*ij*_ for variant *j* of gene *i*. MIRAGE is a probabilistic graphical model that describes a generative process of the variant count data (**Figure 1B**).

**Figure 1.**
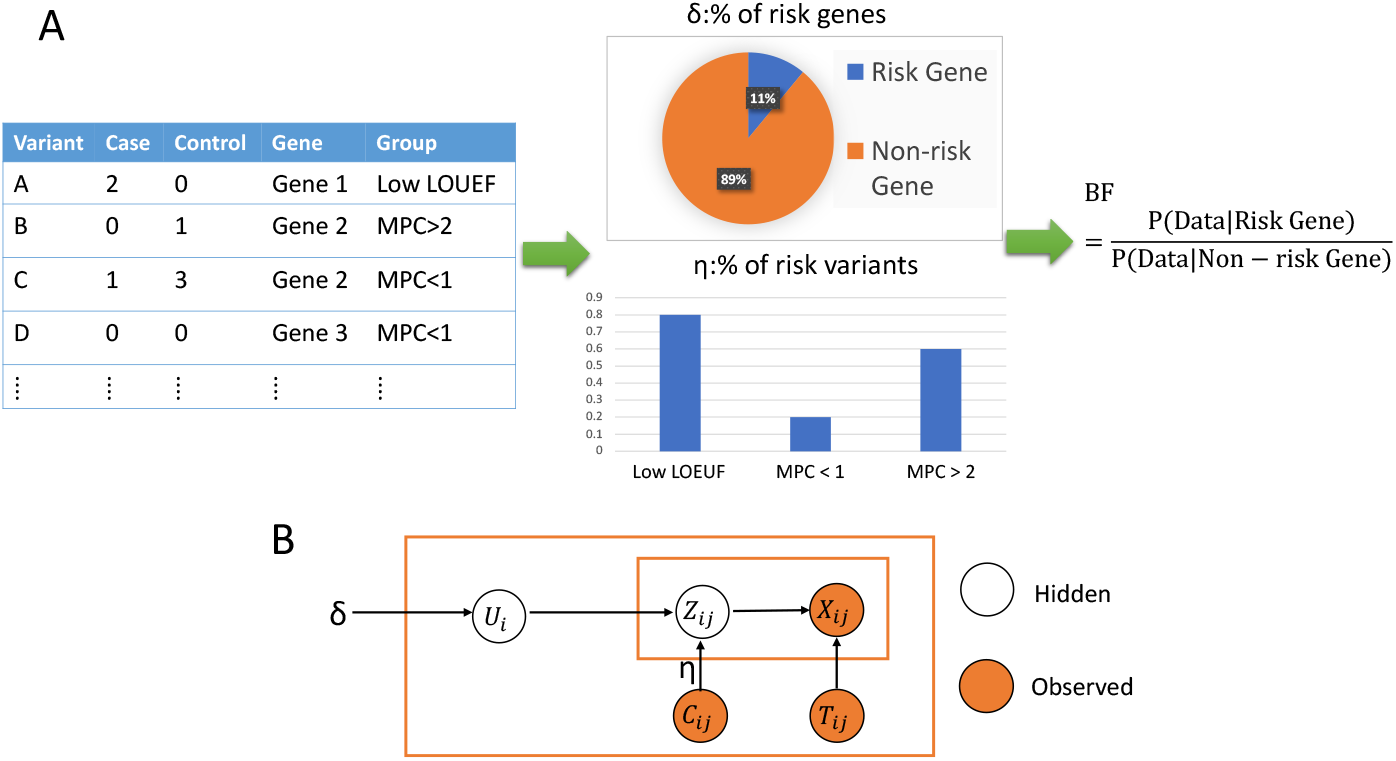
MIRAGE work-flow and model. **(A)** MIRAGE work-flow. The input has information of all rare variants, including case and control counts, associated genes, and the functional groups. MIRAGE estimates the parameters and uses these values to compute the Bayes factor (BF) of all genes. LOEUF: Loss-Of-function Observed/Expected Upper bound Fraction, measures how tolerant a gene is to LoF variants; the lower, the more deleterious. MPC: Missense badness, PolyPhen-2, and Constrain scores; the higher, the more deleterious **(B)** MIRAGE model. See text for definitions of variables and parameters. The outer box corresponds to one gene, and the inner box one variant in a gene. Hidden: unobserved data.

Specifically, we denote *U*_*i*_ the indicator of whether gene *i* is a risk gene. *U*_*i*_ is a Bernoulli random variable with mean *δ*. We denote *Z*_*ij*_ the hidden indicator of whether variant *j* of gene *i* is a risk variant. The distribution of *Z*_*ij*_ depends on the variant category *c*_*ij*_ and the gene indicator *U*_*i*_. When *U*_*i*_ = 0 (non-risk gene), none of gene *i*’s variants would be risk variant, so *Z*_*ij*_ = 0 for all *j*. When *U*_*i*_ = 1 (risk gene), *Z*_*ij*_ would depend on the variant category *c*_*ij*_. We denote *η*_*c*_ the proportion of risk variants in the variant category *c*. Then *Z*_*ij*_ follows Bernoulli distribution with mean 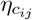.

The allele counts of a variant in cases and controls follow Poisson distributions. Their rates depend on the status of whether a variant is a risk variant. We denote *q*_*ij*_ the allele frequency of variant *j* in the controls. If *j* is a non-risk variant (*Z*_*ij*_ = 0), its allele frequency in cases would also be *q*_*ij*_. So we have:

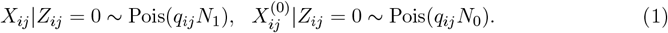

If *j* is a risk variant (*Z*_*ij*_ = 1), its allele frequency in cases would generally be elevated. Let *γ*_*ij*_ be the fold increase of allele frequency. It can be interpreted as the relative risk of variant *j*, as shown in^33^. So we have:

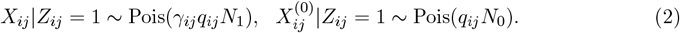

It is generally difficult to estimate *γ*_*ij*_ for individual rare variants, so as in TADA^33^, we treat *γ*_*ij*_ as a random variable following 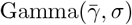. The hyper-parameter 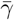 is the prior mean of relative risk of risk variants, and *σ* is the dispersion parameter. These hyperparameters need to be provided by the users, e.g. by estimating them from the data^35^, though in practice, we found that the results are relatively robust to the exact values.

We note that *q*_*ij*_ is a nuisance parameter of no primary interest. So we take advantage of the property of Poisson distribution that the conditional Poisson random variable follows Binomial distribution. This allows us to eliminate *q*_*ij*_:

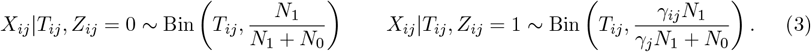

We marginalize *γ*_*ij*_ in evaluating the probability of allele counts for risk variants:

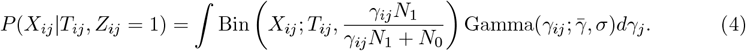

We are now ready to describe the likelihood function and our inference procedure. We denote **X** as the data of allele counts in cases of all variants in all genes. Similarly, we denote **T** the data of allele counts in cases and controls combined of all variants in all genes. We also denote *C* as the set of variant annotations *c*_*ij*_’s. Our primary parameters of interest are *δ*, the proportion of risk genes, and *η*, the vector of *η*_*c*_’s for all variant categories. The likelihood function is given by:

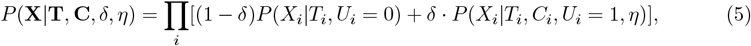

where *X*_*i*_, *T*_*i*_, *C*_*i*_ are the relevant data of all variants in the gene *i*. The first probability term in the equation is the likelihood of a non-risk gene, and is simply given by:

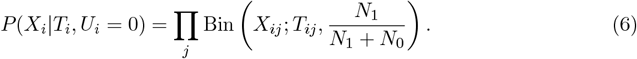

The second probability term is the likelihood of a risk gene:

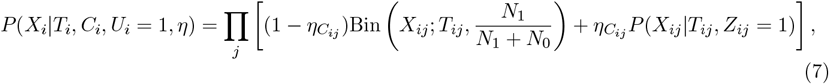

where *P* (*X*_*ij*_|*T*_*ij*_, *Z*_*ij*_ = 1) is given by Equation (4). We note that in computing the likelihood of a single gene, we ignore potential linkage disequilibrium and assume all variants are independent. Since we focus on variants with AF *<* 0.1%, this assumption is generally valid. In practice, we can also perform LD pruning to create independent set of variants.

Given the likelihood function involving latent variables *U*_*i*_’s and *Z*_*ij*_’s, we derive the Expectation-Maximization (EM) algorithm to estimate the parameters *δ* and *η* (see EM Algorithm in the Supplements), with their initial values being randomly chosen. We note that the update rules of the parameters in the M-step have simple, closed forms.

Given the MLE 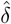 and 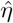, we can determine the Bayes factor of a gene *i, B*_*i*_, and its posterior probability of being a risk gene, PP_*i*_, as:

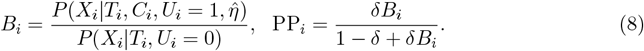

From Equations 6 and 7, it is easy to show that *B*_*i*_ can be related to the evidence at the single variant level:

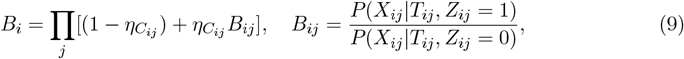

where *B*_*ij*_ is the BF of variant *j* of gene *i*. From this equation, one can see that the more deleterious variant categories with larger values of *η*_*c*_ will contribute more to the gene level evidence. We also note that the log-BF of a gene log *B*_*i*_ can be partitioned as the sum of contributions of each variant, 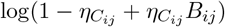. This partition is used when we assessed the contribution of individual variants, or variant groups, to the evidence of a gene in real data analysis. Once we determine the BFs and posterior probabilities of all genes, we control for multiple testing by performing Bayesian FDR control^33^.

### 2.2 Simulation procedure

We simulated the variant counts of a set of genes, 1,000 in our simulations, under case-control data with sample sizes *N*_1_ = *N*_0_ = 3, 000. We assumed each gene has a mixture of variants in different categories, with fixed proportions of variant categories in each gene. We used three categories mimicking LoF, deleterious missense variants and the rest. The proportions of variants in these three categories were 10%, 30% and 60%, respectively. Our simulation started with sampling the risk status for gene *i, U*_*i*_ ∼ Bernoulli(*δ*). When *U*_*i*_ = 0, all variants would be non-risk variants. When *U*_*i*_ = 1, we sampled the risk variant status *Z*_*ij*_ for each variant *j* of gene *i*. For a variant *j* in a category *c*, its probability of being a risk variant *Z*_*ij*_ ∼ Bernoulli(*η*_*c*_). We set *η*_*c*_ 0.5, 0.2 and 0.05 for the three categories, respectively. For a risk variant *Z*_*ij*_ = 1, we sampled the relative risk 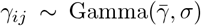. Both 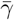 and *σ* were set as user-specified parameters. In the simulations, we used 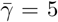 for the first category of variants and 3 for the other two categories, and used *σ* = 1.

Having sampled the status of risk variants and their relative risk, we sampled the variant counts. First, we sampled the allele frequency *q*_*ij*_ from a Beta distribution. If *Z*_*ij*_ = 0, we sampled from Beta(*α*_0_, *β*_0_). We set *α*_0_ = 0.1, *β*_0_ = 1000 in our simulations. If *Z*_*ij*_ = 1, we assumed variants would be even rarer, so we sampled from Beta(*α, β*), where *α* = 0.1, *β* = 2000 (so mean AF is two times lower than non-risk variants). Now we sample the genotype of each individual in cases and in controls. In controls, the genotype of a variant *j* in gene *i* of a subject would follow Bernoulli distribution with mean *q*_*ij*_; and in cases, the genotype would follow Bernoulli distribution with mean *γ*_*ij*_*q*_*ij*_. We note that we sampled the genotype of each variant independently, assuming no LD between variants. From these genotype data, we can collapse variant counts in cases and in controls, which would be used by MIRAGE and other tests.

In additional simulations (**Figure S2**), we varied the number of variants per gene. We sampled the variant number uniformly from 50 to 500 in every gene. The rest of simulations was the same as before.

### 2.3 WES data of ASD

Whole exome sequencing (WES) data of autism were obtained from Autism Sequencing Consortium (ASC) and published in Fu et al^36^. Specifically we aggregated inherited variants from ASC B14, ASC B15-16, SPARK Pilot and SPARK main freeze (WES1), resulting in a total of 14,578 ASD probands and 5,391 unaffected siblings. ASC B14 contains ASC samples and SSC (Simons Simplex Collection), contributing 7,291 probands (6,026 male and 1,265 female) and 2,348 silbings (1,158 male and 1,190 female). ASC B15-16 has 279 probands (223 male and 56 female), and 11 siblings (6 male and 5 female). SPARK main freeze has 6,543 probands (5,219 male and 1,324 female) and 3,032 siblings (1,554 male and 1,478 female). SPARK Pilot has 465 probands (376 male and 89 female) only. To evaluate transmitted/untransmitted variants, dataset specific filters were applied, such as different VQSLOD thresholds. Additional filters were applied to ASC B14 to standardize the samples sequenced across the past several years. After applying these filters, transmitted/untransmitted alleles were called and annotations were produced. We note that the transmitted and untransmitted counts reflect the total number of transmission events, rather than the number of individuals or the specific familial configurations. For instance, we do not distinguish between a variant transmitted once each in two separate families, and a variant transmitted from both heterozygous parents to a single homozygous alternate child in another family. In both cases, the tally would increase by two. More details on the variant calling, variant compilation, quality control can be found in Fu et al^36^, and variant annotations in Satterstrom et al^37^. Synonymous variants and variants with allele frequency *>* 0.1% are filtered.

### 2.4 Applying MIRAGE to ASD data

We annotated variants using both loss-of-function (LoF) and missense annotations. For LoF variants, we applied two commonly used metrics: the probability of loss-of-function intolerance (pLI)^38,39^ and the loss-of-function observed/expected upper bound fraction (LOEUF)^40^, which quantify gene-level intolerance to LoF mutations. For missense variants, we used two approaches: the missense badness, PolyPhen-2, and constraint (MPC) score^41^, and AlphaMis-sense, a deep learning-based pathogenicity predictor^42^.

Based on these annotations, we defined variant groups as follows. For LOEUF: decile 1 (low LOEUF, most intolerant), deciles 2–3 (med LOEUF, moderately intolerant), and deciles *>*3 (high LOEUF, tolerant). For pLI: pLI ≥ 0.995 (high), 0.995 *>* pLI ≥ 0.5 (med), and pLI *<* 0.5 (low). For MPC: MPC ≥ 2 (high), 2 *>* MPC ≥ 1 (med), and MPC *<* 1 (low). For AlphaMissense: likely pathogenic, ambiguous, and likely benign.

All genes in the genome were used to run MIRAGE. The hyperprior parameter for relative risk, 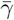, is set at 6 for LoF and 3 for missense variant sets. In the EM algorithm for estimating the model parameter *δ*, we randomly chose initial values, and the algorithm converges if the change of parameter estimates in two iterations is less than 10^−5^. Once these parameters are estimated, their values are assumed to be known, and are used in calculating BF of each gene.

After running MIRAGE on all genes, we checked the LD pattern of the variants supporting all genes at PP *>* 0.5. We defined supporting variants as those with logBF *>* 1. We computed pairwise LD of all the supporting variants in a gene, one gene at a time. From these results, only *LMNB1* and *SV2B* have variants in LD (**Figure S6**). We then chose the variants with the highest logBF, and filtered out the ones in LD. The BFs of the genes would then be updated.

The results of MIRAGE of individual genes were displayed using the lollipop plots. They were generated by trackViewer^43^.

### 2.5 Other programs for rare variant analysis

We used statistical software R (R version 4.4.0) to run other programs, using the same settings in both simulations and real data analyses.

For burden test of a set of variants within a gene, we collapsed the total variant count in cases and in controls, then compared the difference of the burden between the two groups using Fisher’s exact test. The R package *AssotesteR* was employed for CMC and ASUM analyses. For CMC, three MAF cutoffs, i.e. *MAF <* 1*/*3000, 1*/*3000 *< MAF <* 5*/*3000, 5*/*3000 *< MAF <* 20*/*3000, *MAF >* 20*/*3000 were applied to partition the variants. We performed 100 permutations for both CMC and ASUM to obtain p-values, as increasing the number of permutations did not significantly impact the results. R package *SKAT* (version 2.2.5) was used to perform SKAT-O without covariates.

For the ACAT analysis of a gene, we first tested the difference of case and control counts for each variant using Fisher’s exact test. Variants with minor allele counts below 10 were collapsed for testing, following the approach outlined by Liu et al^23^. We then aggregated the p values of all variants within a variant group, using the *ACAT* function from the R package *ACAT* (version 0.91), available at https://github.com/yaowuliu/ACAT. Finally, we applied ACAT to group level p values to get the p value of a gene.

In the analysis of ASD data, we also ran the standard single variant analysis on 1,539,388 variants. This test compares transmitted and non-transmitted variant counts using the binomial test (with the success probability 0.5). As another gene level test, we consider the minimum p value of all variants in a gene. The minimum p values, however, are not calibrated. Under the assumption that all variants are independent, the null distribution of minimum p values follows the Beta distribution, Beta(1, *n*) where *n* is the number of variants in a gene. We thus corrected the minimum p values of all genes using these Beta distributions. We noticed that the resulting empirical p values for both single variant test and minimum-p test are deflated (Supplementary **Figure S5**). This probably reflects that there are a large number of singleton variants (a single total variant count), resulting in p value = 1 from the binomial test. In other words, the p values of most null variants are not uniformly distributed.

### 2.6 Gene set enrichment analysis

MIRAGE identified 18 genes with PP *>* 0.5. We thus initially selected the top 18 genes as identified by MIRAGE, burden tests, and ACAT. Additionally, a comprehensive set comprising all 16,469 genes analyzed were used as a baseline for comparative purposes. Our primary objective was to determine whether these genes exhibit enrichment in gene sets associated with ASD. Here, “enrichment” refers to the proportion of overlap between our top-ranked genes and those within various ASD-related gene sets.

The ASD-related gene sets incorporated into our analysis include: (1) known ASD genes from literature: genes from the SFARI database^44^, including only categories 1 and 2, totaling 769 genes; (2) genes identified in *de novo* mutation studies, TADA^36^ (TADA FDR *<* 0.05, comprising 146 genes); (3) genes associated with Intellectual Disability (ID)^45^, encompassing 252 genes in total; (4) Schizophrenia (SCZ) risk genes^46^, with a total of 1,796 genes; (5) genes involved in relevant biological processes, such as the Post-synaptic density (PSD)^46^, comprising 661 genes, and FMRP target genes^46^, totaling 783 genes; (6) evolutionarily constrained genes as identified by RVIS^47^, including 846 genes, alongside Haplo-insufficient (HI) genes^47^, totaling 1,440 genes; and (7) major depressive disorder (MDD)^48^, totalling 296 genes.

To evaluate the statistical significance of the observed differences in gene enrichment between the MIRAGE-identified genes and the baseline set, Fisher’s exact test was employed.

### 2.7 STRING Network analysis of candidate genes

We used the STRING database (version 12.0, https://string-db.org) to construct gene networks for MIRAGE genes with posterior probability (PP) *>* 0.5 and for autism spectrum disorder (ASD) genes^36^. The ASD genes were identified by jointly analyzing rare *de novo* and inherited protein-truncating variants, damaging missense variants, and copy number variants from exome sequencing of 63,237 individuals, using an extended Bayesian framework (TADA^33^) to integrate variant types and inheritance patterns.

To generate the gene network shown in **Figure 4C**, we applied the default STRING settings. As a control, we randomly sampled 100 genes from all genes analyzed by MIRAGE, constructed their networks, and calculated the number of links between these random genes and ASD genes.

We then compared the number of links of MIRAGE genes to ASD genes against that of random genes using a Poisson test. Specifically, we observed 42 links between 18 MIRAGE risk genes and ASD genes. In the random set, we observed 1.46 links per gene. We would then test the significance using Poisson test.

## 3 Results

### 3.1 Overview of MIRAGE

MIRAGE requires summary statistics as the input, in the form of the counts of all rare variants in cases and in controls (**Figure 1A**). In addition, MIRAGE takes functional information of variants as input, assuming variants are assigned into disjoint categories, based on their likely effects, for example, loss-of-function (LoF) variants, and likely deleterious missense variants^41^ (**Figure 1A**). For each gene, MIRAGE assesses how likely its data is generated from a non-risk gene model *M*_0_, vs. a risk gene model *M*_1_. Under *M*_0_, all the variants of a gene are non-risk variants, and their expected frequencies are equal between cases and controls. Under *M*_1_, any variant has a prior probability of being a risk variant, with the probability *η*_*c*_ for the variants in the category *c*. The frequencies of risk variants would be generally different between cases and controls. Thus each variant would contribute some information: an imbalance between case and control counts would provide some support to *M*_1_, and functionally important variants would make larger contributions. The information across all variants in a gene would then be combined by MIRAGE to form the statistical evidence of *M*_1_ vs. *M*_0_, known as Bayes factor (BF).

MIRAGE estimates all the parameters, including the prior probability of being a risk gene, *δ*, and the proportion of risk variants, *η*_*c*_, for all categories, using the entire dataset of all genes (**Figure 1A**). Then for each gene, it assesses its evidence as a risk gene by computing its BF. The BF of a gene can be used to compute its Posterior Probability (PP) of being a risk gene.

Unlike p value, PP has a simple interpretation of the probability of being a risk gene, given all the data we have about the gene. Using the PPs, MIRAGE can also perform multiple testing control, using a Bayesian False Discovery Rate (FDR) approach^49^.

MIRAGE can be formulated as a probabilistic graphical model (**Figure 1B**). Let *U*_*i*_ be an indicator variable of whether the gene *i* is a risk gene (1 if yes and 0 otherwise). Then *U*_*i*_ follows a Bernoulli distribution with probability *δ*. For the *j*-th variant of gene *i*, let *X*_*ij*_ and *T*_*ij*_ be its allele count in cases and the total allele count, respectively. The distribution of these counts depends on whether the variant *j* is risk variant or not, denoted as *Z*_*ij*_. For a non-risk gene (*U*_*i*_ = 0), *Z*_*ij*_ = 0 for all variants. For a risk gene (*U*_*i*_ = 1), *Z*_*ij*_ depends on its variant category, denoted as *C*_*ij*_. If its category is *c*, then *Z*_*ij*_ is a Bernoulli random variable with probability *η*_*c*_. Given *Z*_*ij*_ of a variant, we can model its allele count in cases as a binomial distribution. Assuming equal numbers of cases and controls (see Methods for the general setting), we have:

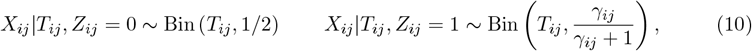

where *γ*_*ij*_ is the relative risk of variant *j*, modeled as a Gamma distribution. We assumed all variants are independent. We think this is justified when one analyzes rare or ultra-rare variants, with MAF below 0.1%. In practice, we can also prune variants in LD. With this probabilistic model, we used an Expectation Maximization (EM) algorithm^50^ to estimate the proportion of risk genes (*δ*), and the proportions of risk variants, *η*_*c*_’s. Once the parameters were estimated, we computed the BF of a gene *i*, as the ratio of the probability of all its variant counts, *X*_*ij*_’s, under *U*_*i*_ = 1 vs. *U*_*i*_ = 0 (see Methods). We note that the log-BF of a gene only depends on the *η*_*c*_ parameters and can be partitioned as the sum of contributions of each variant (see Equation 8 and 9 in Methods). This partition allows us to assess the contribution of individual variants, or variant groups, to the evidence of a gene.

### 3.2 MIRAGE is more powerful in identifying risk genes than existing methods in simulations

We simulated data by using the genetic architecture information from ASD, to compare MIRAGE with existing methods in identifying risk genes from rare variants. To demonstrate the power of MIRAGE even in small samples, we fixed sample sizes at 3,000 cases and 3,000 controls. We simulated data of 1,000 genes, with the proportion of risk genes, *δ*, varying from 0.02, 0.05, 0.1, to 0.2. For simplicity, we assumed every gene has the same number of variants (100 variants per gene), however, the results were similar if we vary the number of variants from 50 to 500 (**Figure S2**). For a risk gene, its variants fall into three functional categories, mimicing benign missense variants (60% of variants), damaging missense variants (30%) and Loss-of-function (LoF) variants (10%), respectively. The proportions of risk variants, *η*_*c*_, are 0.05, 0.2 and 0.5 for the three categories. Once the risk status of a variant is sampled, the allelic counts of the variant in cases and controls would follow Poisson distributions. The ratio of the two Poisson rates is the relative risk of that variant, sampled from Gamma distributions. Based on earlier exome sequencing studies of ASD^33,51^, we set the mean relative risks, 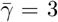 for the first two variant categories and 5 for the last one.

We compared MIRAGE to several variations of burden tests (a baseline Burden test, CMC^16^ and ASUM^52^), SKAT and ACAT. The baseline burden test tested all variants within a gene.

In practice, Burden test is often applied to different categories of variants of a gene separately to increase the power. We thus considered two other versions of burden test as well. The first version tested each of the three variant categories separately, then used the minimum p value of the three, adjusting for three tests. The second combined the three burden p values from the variant categories using Fisher’s method. The results of these two tests in simulations were similar to the baseline Burden test (**Figure S1**), so we considered only the baseline version here.

We compared the performance of the methods in distinguishing risk from non-risk genes, using the ROC curves (*δ* = 0.02 and *δ* = 0.1 in **Figures 2A–2B**, *δ* = 0.05 and *δ* = 0.2 in **Figure S2**).

**Figure 2.**
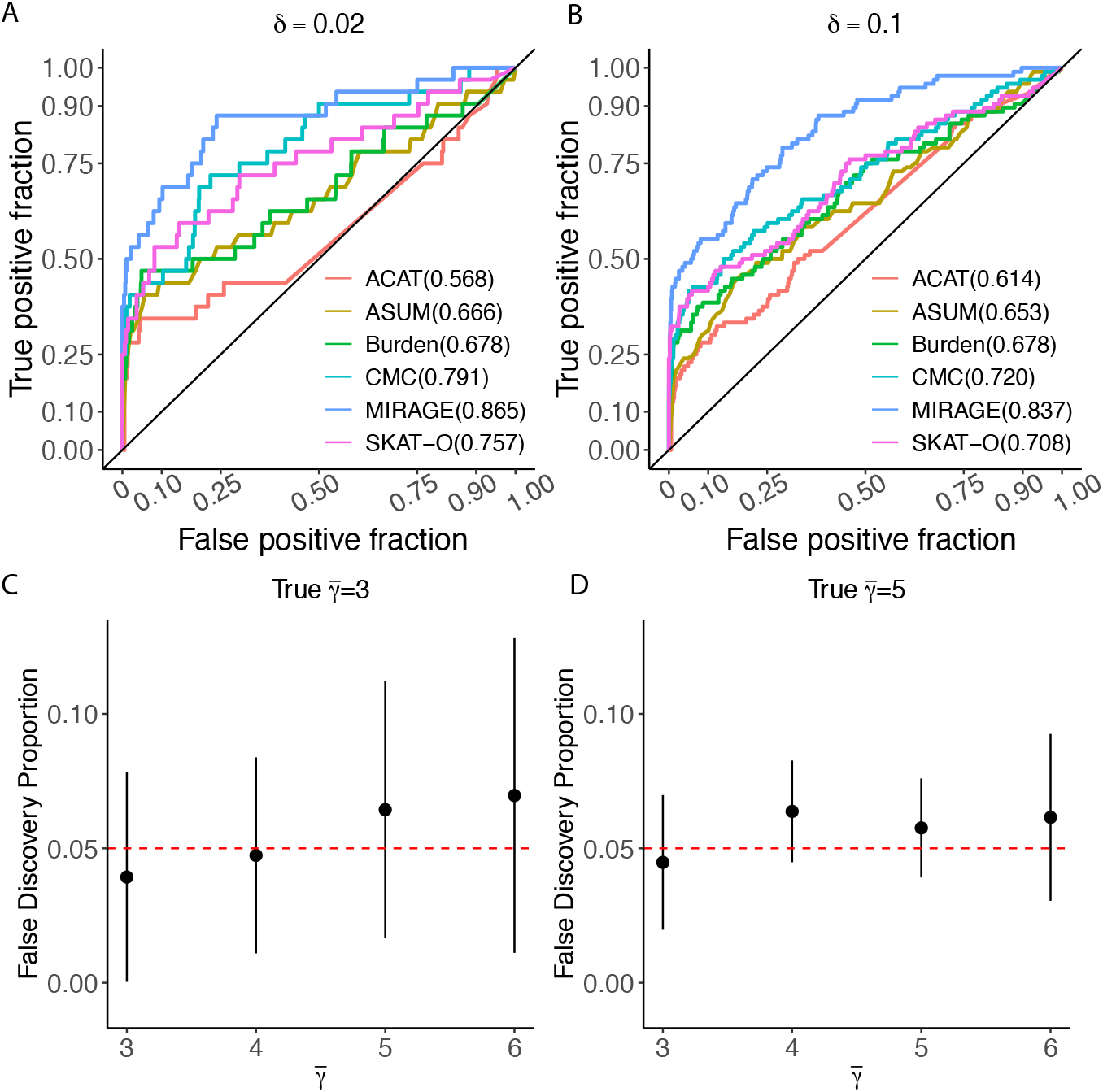
Simulation results. **(A)** ROC curves at *δ* = 0.02 of different methods for classifying risk genes. We simulated 1,000 genes with varying proportion of risk genes. AUC values are shown in the bracket. Solid black reference line is in the diagonal. **(B)** ROC curves at *δ* = 0.1 of different methods for classifying risk genes. **(C)** False discovery rate(FDR) calibration by MIRAGE. We simulated 20 datasets with true 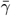 (relative risk) = 3. MIRAGE used 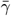 from 3 to 6. The False Discovery Proportion under each specified value of 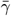 is shown. Red dashed line is the target FDR level (0.05). **(D)** FDR calibration by MIRAGE at 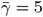

Under all the settings, we found that MIRAGE has Area Under Receiver Operating Characteristic (AUROC) above 80%, substantially outperforming all other methods. To appreciate how much of this difference translates into the difference of power of detecting risk genes, we estimated the number of genes found by each gene at FDR *<* 0.1 (for MIRAGE, we used Bayesian FDR). We found that the power of MIRAGE is the highest in all settings, and ∼ 20% higher than the second best method (SKAT-O) in the simulations (**Figure S3**).

MIRAGE uses a Bayesian approach to controlling FDR. We performed additional simulations to assess if the Bayesian FDR is calibrated, and whether the FDR is sensitive to misspecification of 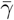, the average relative risk of risk variants. To make simulations simpler, we ran similar simulations as before, but used a common value of 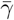 for all three variant categories. In simulations, we used a range of values, from 3 to 6, as 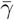. When running MIRAGE, we assumed that the true value of 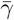 was unknown and ran MIRAGE using the value ranging from 3 to 6. Our results were generally robust to the value of this parameter, and the Bayesian FDR was close to the true False Discovery Proportions (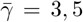 in **Figures 2C–2D**, 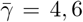 in **Figure S4**).

### 3.3 MIRAGE identifies putative risk genes of ASD

We applied MIRAGE to WES data of 14,578 trios of children affected with ASD and their parents, combining data from Autism Sequencing Consortium and SPARK^36^. We treated transmitted parental alleles as “cases” and non-transmitted ones as “controls”^33^. We considered only rare variants with minor allele frequency (MAF) (both within cohort MAF and gnomad non-neuro allele frequency) below 0.1%, and filtered all synonymous variants. To annotate variants, we used LOEUF for LoF mutations^40^, and MPC scores (Missense badness, PolyPhen-2, and Constrain scores) for missense variants^41^. LOEUF quantifies how much a gene tolerates loss-of-function mutations and is widely used. MPC has been shown to enrich risk variants for ASD in previous studies^37,41^. We included a total of six variant groups defined by the LOEUF decile and MPC score: decile 1 (low LOEUF, the lower, the more deleterious), deciles 2 and 3 (med LOEUF), and deciles greater than 3 (high LOEUF), MPC≥ 2 (high MPC), 2 *>* MPC ≥ 1 (med MPC) and MPC *<* 1 (low MPC).

We first confirmed that the ASD dataset was challenging for current methods. We applied Burden tests and ACAT to all genes. We used two definitions of burden: 1) burden in all LoF variants; and 2) burden in LoF and missense variants at MPC *>* 2. In ACAT, we conducted a binomial test on each variant within a gene, comparing the number of transmitted and nontransmitted variants. The statistics of variants within a gene were then combined using the ACAT method, with the variant weights determined by the MAFs^40^, as described previously^23^. The QQ plots of p values showed that none of these methods detected any signal (**Figure 3A**). As other baseline methods, we also considered the single-variant test, as well as a gene level test using minimum p value of all variants (adjusting for the number of variants). Neither test showed any significant findings (**Figure S5**).

**Figure 3.**
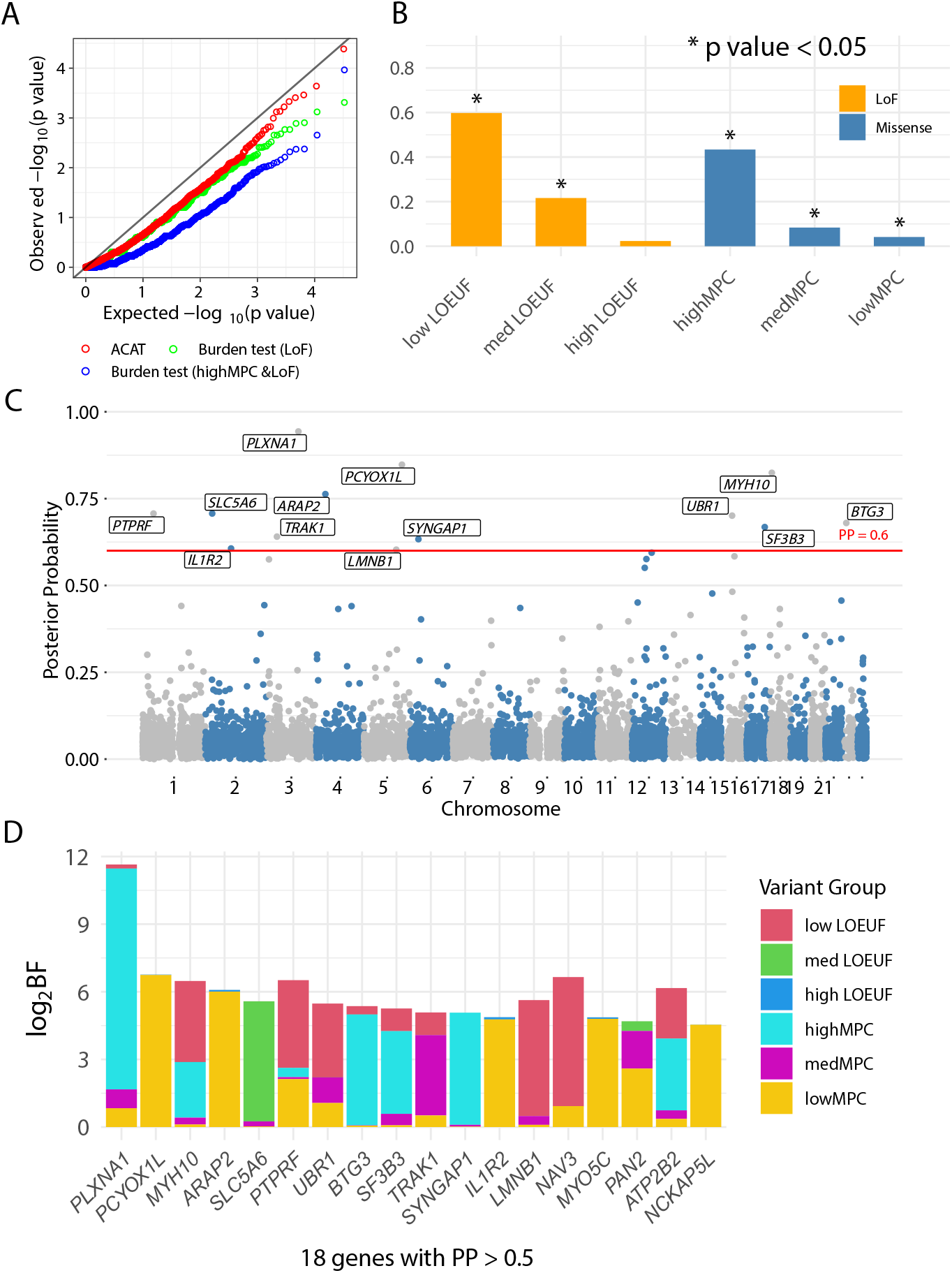
Summary of MIRAGE results from ASD data analysis. **(A)** QQ plot of all genes in the whole genome by ACAT and burden tests. **(B)** The proportion of risk variants in 6 variant categories. LoF categories: LOEUF (Loss-Of-function Observed/Expected Upper bound Fraction) decile = 1 (low), LOEUF decile = 2–3 (med), LOEUF decile = 4–10 (high). Missense categories: MPC (Missense badness, PolyPhen-2, and Constrain scores) ≥ 2 (high), 2 *>* MPC ≥ 1 (med), MPC *<* 1 (low). **(C)** Manhattan plot of posterior probabilities (PP) for all gene analyzed. Genes with PP *>* 0.6 are labeled. **(D)** The log_2_BF (BF is defined in **Figure 1A**) of a gene is partitioned into contributions of variants in each group. Shown are results of 18 genes at PP *>* 0.5, genes are ordered by their PPs. The negative log_2_BF of a variant group is truncated to 0.

To run MIRAGE, we used an earlier estimate based on *de novo* mutations that 5% of genes are ASD risk genes^53^. Given that the signals in the transmission data is considerably weaker than *de novo* mutations^36^, we think this estimate is more accurate than the value inferred from transmission data alone. Using this proportion of risk genes, MIRAGE estimated the proportions of risk variants across the six variant categories (**Figure 3B**). The LoF variants in genes with high LOEUF score showed the highest proportions, with ∼60% of variants being risk variants. In the missense variant groups, high MPC had the largest risk variant proportion of 43%, and the last two categories showed much lower proportions. These estimates were generally in line with the expected deleteriousness of these variant annotations.

With the estimated parameters, we calculated BFs and posterior probabilities (PP) for every gene and performed Bayesian FDR control. At PP *>* 0.5, MIRAGE identified 18 putative ASD risk genes (**Figure 3C, Table S1**). We verified that the vast majority of the variants supporting these genes have very low LD (**Figure S6**), and pruning of the few remaining variants in LD have small effects on the results (see Methods). At PP *>* 0.7, MIRAGE identified seven putative ASD risk genes. These results thus supported higher sensitivity of MIRAGE in detecting risk genes than existing rare variant association methods.

To better understand the MIRAGE results, we evaluated the contributions the variant groups to the associations of these genes with ASD. Following earlier work^36^, we partitioned the evidence of each gene, in terms of log_2_BF, into contributions of the six variant groups, in all genes at PP *>* 0.5 (**Figure 3D**). We found that the variant group(s) driving association signals in each gene varied considerably. For instance, *PLXNA1* was predominantly driven by the high MPC group of missense variants, while the association signals for *MYH10* was largely attributed to LoF variants in genes with low LOEUF. In about half of the genes, two or more variant groups made substantial contributions. Overall, the LoF variants from genes with Low/Med LOEUF and the High MPC variant group drove the associations in most of the genes. Together, these results highlighted that MIRAGE was able to effectively combine statistical signals across multiple variant groups to discover risk genes.

Finally, we evaluated the MIRAGE results using different functional annotations, including pLI for LoF variants, and AlphaMissense^42^ for missense variants and pLI for LoF variants, and MPC for missense variants. The results from using these annotations broadly agreed with the results here. The estimated parameters reflected the expected severity of variants in the functional categories, and the discovered genes showed enrichment in ASD-related gene sets (**Figures S13–S16**)

### 3.4 Candidate genes found by MIRAGE are supported by multiple lines of evidence

We evaluated the candidate genes by assessing the enrichment of ASD-related gene sets. We selected the 18 genes at PP *>* 0.5, and for comparison, the same number of top genes by burden tests and ACAT. The ASD-related gene sets include known ASD genes curated by SFARI^44^ and from de novo mutation studies using TADA^36^; risk genes of Intellectual Disability (ID)^45^ and Schizophrenia (SCZ)^46^; relevant biological processes including Post-synaptic density (PSD)^46^ and FMRP target genes^46^; evolutionarily constrained genes from RVIS^47^ Haplo-insufficient (HI) genes^47^ and major depressive disorder (MDD)^48^. We note that the SFARI and TADA genes were derived from independent datasets. While some samples in our dataset were included in earlier studies, the transmission data were not used previously. We found that the top MIRAGE genes were enriched in most ASD related gene sets (**Figure 4A**). For example, ∼ 35% of MIRAGE candidate genes were likely ASD genes by SFARI (*p* = 6.7 *×* 10^−5^, Fisher’s Exact Test). The top genes from other methods showed lower or no enrichment. These enrichment results thus strongly supported the likely roles of the candidate genes in ASD.

**Figure 4.**
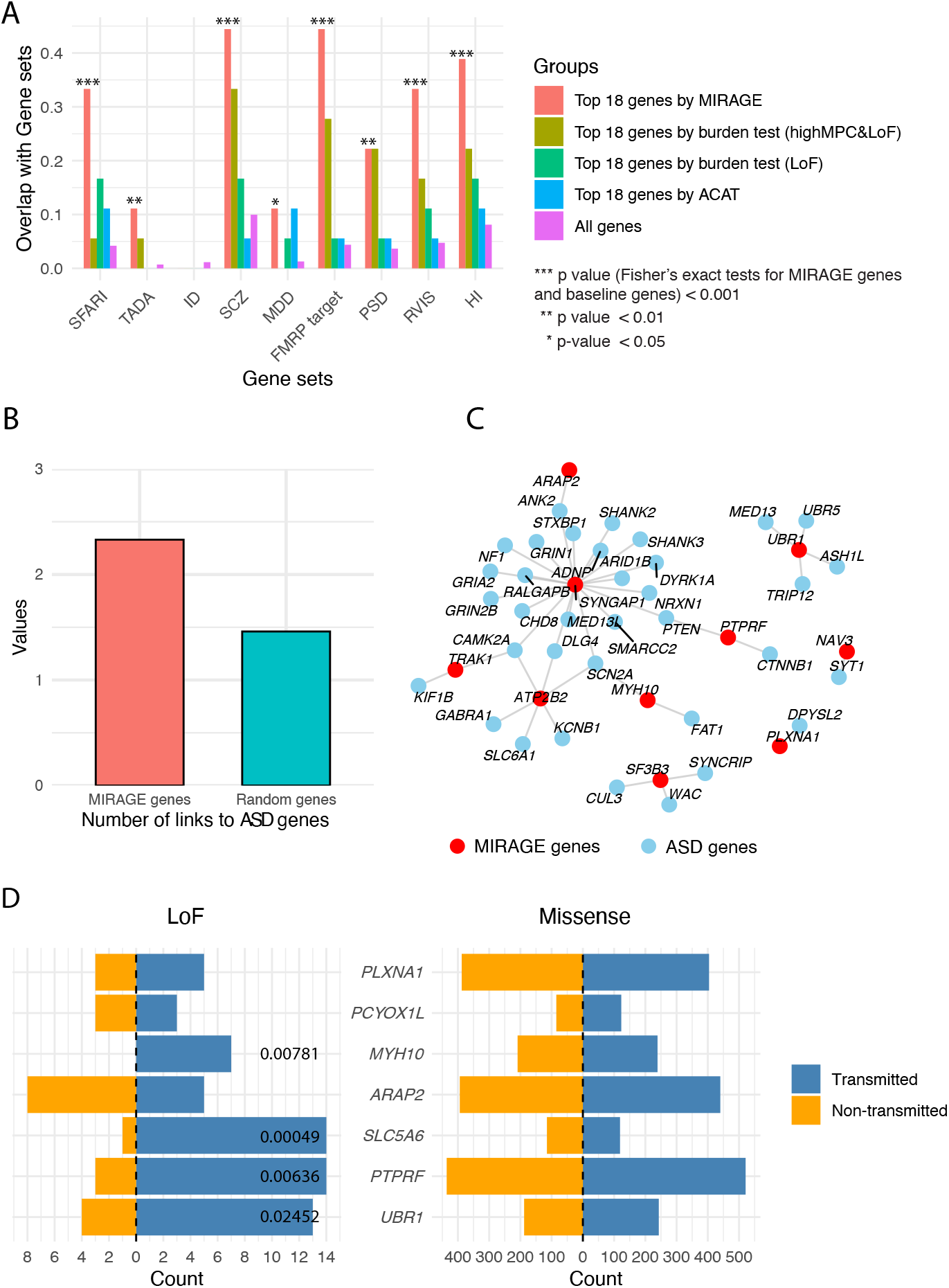
Evaluation of candidate genes from MIRAGE. **(A)** Enrichment of ASD-related gene sets in the top 18 genes found by MIRAGE, burden tests and ACAT. Enrichment test was based on Fisher’s exact test. SFARI genes: Categories 1 and 2. TADA high confidence genes: FDR *<* 0.05. ‘All genes’ refer to the entire set of genes analyzed, as a baseline. **(B)** Number of links from MIRAGE genes to TADA genes. **(C)** Gene interaction network for MIRAGE genes and TADA genes, only genes with connections are shown here. **(D)** The transmitted and non-transmitted counts of LoF (left) and missense (right) variants, for 7 genes with MIRAGE PP *>* 0.7. Numbers near the bars are *p*-values testing transmission disequilibrium, using the binomial test (only shown *p <* 0.05).

We next examined the function and plausibility of the top 7 genes using the more stringent cutoff, PP *>* 0.7 (Table 1). 6 out of 7 genes were annotated by one or more ASD-related gene sets. Among the six genes, *PLXNA1* were found by in earlier ASD studies^34,36^. *MYH10* was a ASD risk gene according to SFARI (score 2, Strong candidate). It was supported by multiple *de novo* mutation studies^54–57^. *PTPRF* and *ARAP2* have known or related functions in neurodevelopment and/or other neuropsychiatric disorders. For example, both of them have been implicated in SCZ. We discussed the literature support of these genes in Discussion.

**Table 1.**
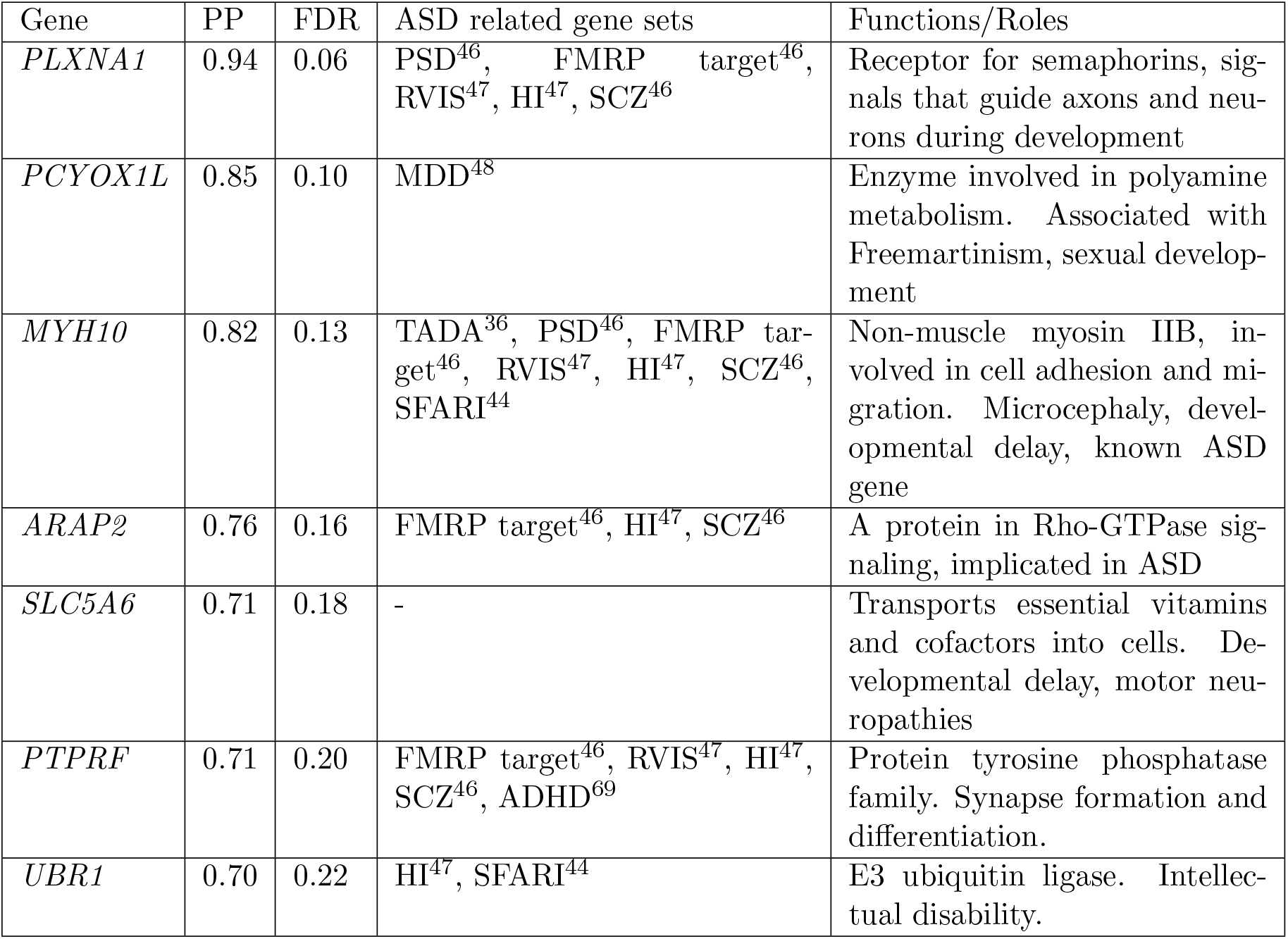
Genes identified by MIRAGE (posterior probability *>* 0.7). PP (Posterior Probability). FDR (False Discovery Rate). The dash (-) indicates no data.

| Gene | PP | FDR | ASD related gene sets | Functions/Roles |
| --- | --- | --- | --- | --- |
| *PLXNA1* | 0.94 | 0.06 | PSD<sup>46</sup>, FMRP target<sup>46</sup>, RVIS<sup>47</sup>, HI<sup>47</sup>, SCZ<sup>46</sup> | Receptor for semaphorins, signals that guide axons and neurons during development |
| *PCYOX1L* | 0.85 | 0.10 | MDD<sup>48</sup> | Enzyme involved in polyamine metabolism. Associated with Freemartinism, sexual development |
| *MYH10* | 0.82 | 0.13 | TADA<sup>36</sup>, PSD<sup>46</sup>, FMRP target<sup>46</sup>, RVIS<sup>47</sup>, HI<sup>47</sup>, SCZ<sup>46</sup>, SFARI<sup>44</sup> | Non-muscle myosin IIB, involved in cell adhesion and migration. Microcephaly, developmental delay, known ASD gene |
| *ARAP2* | 0.76 | 0.16 | FMRP target<sup>46</sup>, HI<sup>47</sup>, SCZ<sup>46</sup> | A protein in Rho-GTPase signaling, implicated in ASD |
| *SLC5A6* | 0.71 | 0.18 | - | Transports essential vitamins and cofactors into cells. Developmental delay, motor neuropathies |
| *PTPRF* | 0.71 | 0.20 | FMRP target<sup>46</sup>, RVIS<sup>47</sup>, HI<sup>47</sup>, SCZ<sup>46</sup>, ADHD<sup>69</sup> | Protein tyrosine phosphatase family. Synapse formation and differentiation. |
| *UBR1* | 0.70 | 0.22 | HI<sup>47</sup>, SFARI<sup>44</sup> | E3 ubiquitin ligase. Intellectual disability. |

To further validate and characterize the functions of the candidate genes, we used STRING^58^, a widely used resource for gene interactions, to assess the connectivity of our top genes with known ASD genes. Our rationale is that true risk genes would tend to be connected with known risk genes through physical or other forms of functional interactions. We assessed the top 18 genes found by MIRAGE at PP *>* 0.5, and compared the connectivity with 100 randomly chosen genes. We found that a MIRAGE gene is connected to 2.33 ASD genes^36^ on average, significantly higher than random genes (1.46) (p = 0.004, Poisson test, **Figure 4B**). These connections provided important clues of how the identified risk genes from MIRAGE may affect the ASD risk (**Figure 4C**). For example, *PTPRF* (PP = 0.71) encodes a receptor-type protein tyrosine phosphatase. It is linked to known ASD genes, *PTPN* and *CTNNB1*, two genes important for PI3K signaling and Wnt signaling, respectively. These connections thus suggested that *PTPRF* may act on ASD by modulating the PI3K and/or WNT signaling, two pathways important for ASD^59–62^. As another example, *UBR1* (PP = 0.7) is linked to several ASD genes, including *UBR5* and *TRIP12*. All three genes have functions in ubiquitination^63–65^ and regulation of protein turnover^66–68^.

We evaluated the statistical support of a few genes in more details. In MIRAGE, the evidence of a gene, log_2_BF, is the sum of log_2_BF of individual variants, allowing us to quantify the contribution of individual variants. For this analysis, we plotted the transmitted and nontransmitted counts of each variant, and its log_2_BF. We highlighted here the result of *ARAP2*, a signaling gene. Its association with ASD was largely driven by two missense variants that are 4bp away (**Figure 5**). The two variants are located in the PH3 domain, a domain involved in binding phosphatidylinositol phosphates, an important class of signaling molecules. In other top genes, the association signals tend to be more diffuse, with multiple supporting variants (**Figures S7–S12**).

**Figure 5.**
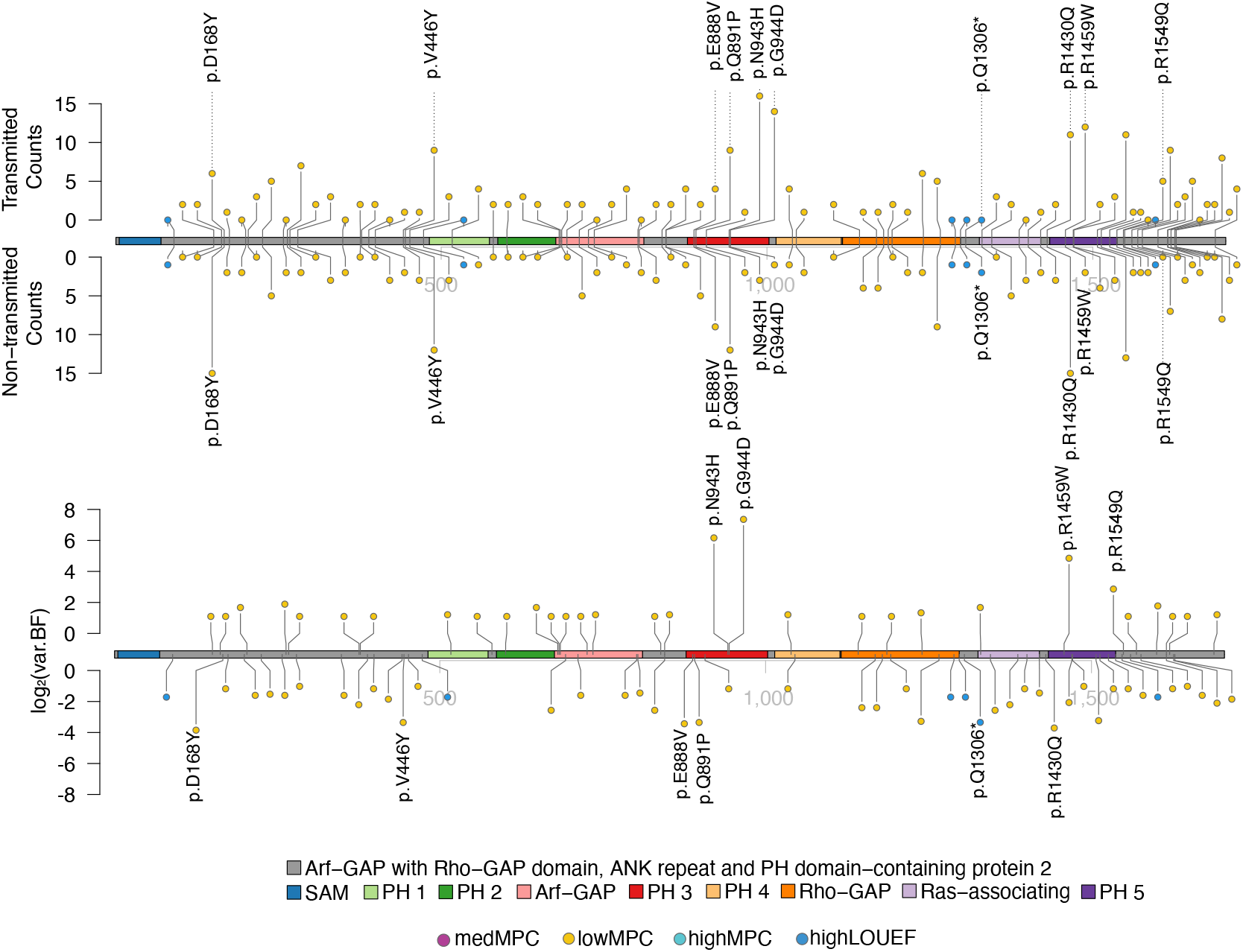
The lollipop plot of *ARAP2*. Top: the transmission and non-transmission counts of each variant. Bottom: the evidence (log_2_BF) of each variant. Only informative variants, defined as BF *>* 2 or BF*<* 0.5 (BF is defined in **Figure 1A**), were plotted. The variants with BF *>* 3 and BF *<* 0.1 were labeled. LOEUF (Loss-Of-function Observed/Expected Upper bound Fraction). MPC (Missense badness, PolyPhen-2, and Constrain scores).

For comparison, we investigated transmission disequilibrium for the LoF and missense variants, each as a group, of the top seven genes. No apparent burden was observed in the missense variants (**Figure 4D**, right). We note that MIRAGE is able to detect signals in the missense variants even when there is no overall burden between transmitted and non-transmitted variants, e.g. for *PLXNA1* and *ARAP2* (**Figure 3D** and **Figure 4D**). While disequilibrium was observed in the LoF group in four genes (at *p <* 0.05) (**Figure 4D**, left), the p values would not pass the genomewide multiple testing threshold.

These results together supported the key rationales of MIRAGE. First, by modeling the heterogeneity of variant effects, MIRAGE allows a small number of variants to drive the results, and is also able to leverage the collective signal across many variants. This benefit is clear in the case of missense variants, where burden test failed to detect any burden (**Figure 4D**, right). Secondly, by borrowing information across genes, MIRAGE is able to learn to put more emphasis on more deleterious variant groups. This allows MIRAGE to extract statistical signals from relatively modest burden of LoF variants (**Figure 4D**, left).

## 4 Discussion

We proposed a Bayesian method, MIRAGE, for rare variant association test. MIRAGE addresses two key limitations of current methods. By treating all variants as a mixture of risk and non-risk variants, it better models the heterogeneity of variant effects, particularly the sparsity of risk variants. Furthermore, it provides a rigorous framework to leverage functional annotations of variants in identifying risk genes. Simulations confirmed the effectiveness of our method. In application to a WES dataset of ASD, while existing methods failed to detect significant associations, MIRAGE identified a number of candidate genes. The top genes were highly enriched with ASD-related gene sets. Most of the six candidate genes, at PP *>* 0.7, are either reported ASD genes, or have functions in related phenotypes or neurodevelopment, representing plausible ASD risk genes.

How to effectively analyze rare variants is a key challenge of the field. The success of MIRAGE in the study of ASD offered some general lessons. First, the effects of rare variants are likely heterogeneous, and this is better captured by a sparse model where most rare variants have no effects on disease risks. One can see this point from our estimated fractions of risk variants, especially in missense variants (**Figure 3B**), and from the analysis of individual genes (**Figure 5, Figures S7–S12**). Secondly, using external information of variants is critical to improve the signal to noise ratio. Indeed, annotating function of variants is an active area of research, and some recent methods, e.g. those based on deep learning^70–72^, may further boost the power of MIRAGE.

MIRAGE is related to some Bayesian statistical methods for analyzing rare variants^73–80^. These methods typically treat the effect sizes of variants as random variables, and some methods explicitly capture the dependency of the effect sizes on variant annotations. For example, in MiST, the prior mean of the effect sizes is a linear function of variant annotations^76^. In a method that generalizes SKAT, the prior variance of effect size is modeled as a function of variant annotations^77^. These methods, however, are not widely used in practice, likely due to several limitations. The effect sizes are often modeled as continuous random variables, similar to SKAT. As we have explained and demonstrated, the risk variants are likely sparse, especially for missense variants. So a mixture approach with the majority of variants as non-risk variants is likely more powerful. Secondly, the existing Bayesian methods analyze one gene at a time. We think it is difficult to learn the parameters about functional annotations from a single gene. For instance, a gene may have only one or few LoF variants, and there is simply not enough information to learn that LoF variants are important. Existing methods may deal with this issue by allowing user-specified weights. We think a better approach is to use Empirical Bayes to borrow information across many genes to estimate the hyperparameters. Lastly, some of these Bayesian methods rely on computationally expensive procedures and are thus difficult to use in practice to analyze thousands of genes. We think these limitations are largely addressed by MIRAGE, through its mixture model of variants, the EM algorithm to estimate hyperparameters from all genes, and the simple binomial distributions for variant counts.

When applying MIRAGE to trio-sequencing data, it is related to TDT on rare variants.

Several methods have been proposed to do rare variant analysis on trio- and family-sequencing data, such as RV-TDT^31^, PedGen^81^, and rare variant Family-based association test (FBAT)^82,83^. These methods are often extensions of burden test, SKAT or other p value aggregation methods, while accounting for family structure. For example, in De et al^82^ and in RV-TDT, the TDT or FBAT statistics were modified by replacing allelic counts of a single variant with genetic burden across multiple variants. In other methods, one may first obtain family-based test statistic (e.g. FBAT) of a single variant, then aggregate over variants using SKAT, or p value aggregation methods^83^. Regardless of the specific approaches, all these methods are closely related to the existing approaches for rare variant analysis, and are thus subject to similar weakness as reviewed earlier.

We discussed in more details the functions of the identified ASD candidate genes. *PTPRF* is a member of protein tyrosine phosphatase (PTP) family. The subfamily containing *PTPRF* plays an important role in neural function such as dendrite and axon growth and guidance, synapse formation and differentiation^84^. A related protein, *PTPRF* interacting protein alpha 1, is a strong ASD risk gene by SFARI^44^. *ARAP2* is an adapter protein in Rho GTPase signaling. A number of studies have linked Rho GTPase signaling to the pathogenesis of ASD^85^. The gene family of *ARAP2* (Arf GAP) are important regulators of the Actin Cytoskeleton, and have been implicated in cell adhesion, motility and neurite outgrowth^86^. *SLC5A6*, which transports biotin and pantothenic acid to the brain, is implicated in brain development and developmental delay^87–89^, while *UBR1* is associated with Johanson–Blizzard syndrome, a disorder characterized by cognitive impairment^90^. We presented the supporting evidence from literature of all genes at PP *>* 0.5 in **Table S2**.

Despite the strong support of our candidate genes from gene set enrichment analysis and literature evidence, it is important to experimentally validate the functions of the identified ASD risk genes in future studies. This may involve CRISPR perturbations of these genes in appropriate cellular models^91^, followed by assessment of molecular and cellular phenotypes, including gene expression, neuronal morphology and synaptic activity.

We briefly commented here about how to run MIRAGE. First, we note that the main parameters of MIRAGE are *δ*, the proportion of risk genes, and *η*_*c*_, the proportion of risk variants in each functional category. While MIRAGE is able to estimate all the parameters using EM, in practice, it may be better to specify *δ*, if some approximate value can be provided. The model may have low identifiability when the signal is weak, in other words, different values of *δ* may give similar likelihood. Next, MIRAGE results depend on functional annotations used. What is the best annotation in a particular dataset may be unclear a priori. For example, while AlphaMissense works well in other contexts, it does not seem to work as well as MPC scores in our ASD dataset (**Figure 4A** vs. (**Figures S13–S14**)). Choosing what annotations to use may be an important consideration in applying MIRAGE.

MIRAGE can be further developed in several directions. First, MIRAGE was designed for case-control, or transmission studies. Extending it to quantitative traits would greatly broaden its applications. Secondly, MIRAGE does not accommodate sample covariates in analysis, such as age, gender, and population ancestry. Population stratification is of particular concern as it may lead to false positive findings. Incorporating such covariates is thus an important next step. Lastly, MIRAGE currently supports only disjoint functional groups as annotations. This simplifies the mathematical model, but restricts the number and types of annotations one may use. A future direction is to have more flexible models of the prior probabilities of variants. This type of prior has been used successfully in GWAS^92,93^ and in studies of *de novo* mutations^51^.

## Data and code availability

MIRAGE is available at https://xinhe-lab.github.io/mirage/.

## Acknowledgements

This work was supported by the National Institutes of Health (NIH) under grants R01MH110531, R01MH106575, and R01HL163523 (to X.H.). We thank other members of He’s groups for helpful comments on the work and the manuscript.

## Author Contributions

Conceptualization, X.H.; Methodology, S.H., X.H.; Software, G.W., X.S., S.H.; Data Analysis, S.H., X.S., L.S., X.X.; Data Curation, S.H., X.S., L.S., F.K.S., G.W., L.L., N.K., S.Z., Y.L., W.S., H.N., J.B.; Writing, S.H., X.S., X.H.; Supervision, X.H., J.B.; Project Administration, X.H.; Funding Acquisition, X.H..

## Web Resources

ASC data: dbGaP (accession ID: phs000298). SPARK data: https://www.sfari.org/resource/spark/.

## Supplementary Information

## Supplemental Figures

**Figure S1.**
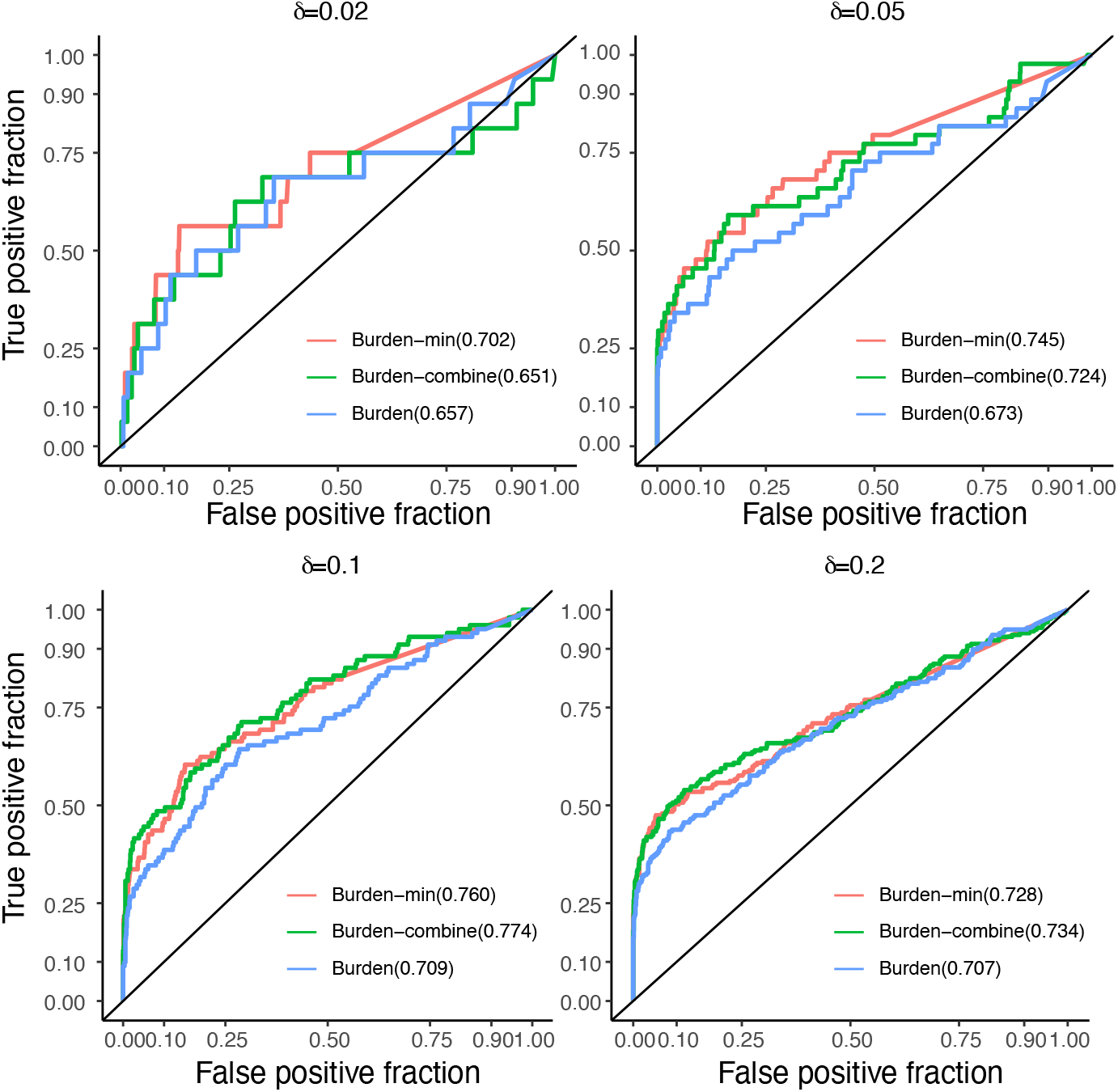
ROCs of several variations of burden test. Burden (basic version): calculates the p value for all variants collectively. Burden-min: calculates the p value for each variant category separately, applies Bonferroni correction to adjust the resulting three p values for each gene, and selects the minimum adjusted p value. Burden-combine: combines the original p values from the burden test across the three categories using Fisher’s method.

**Figure S2.**
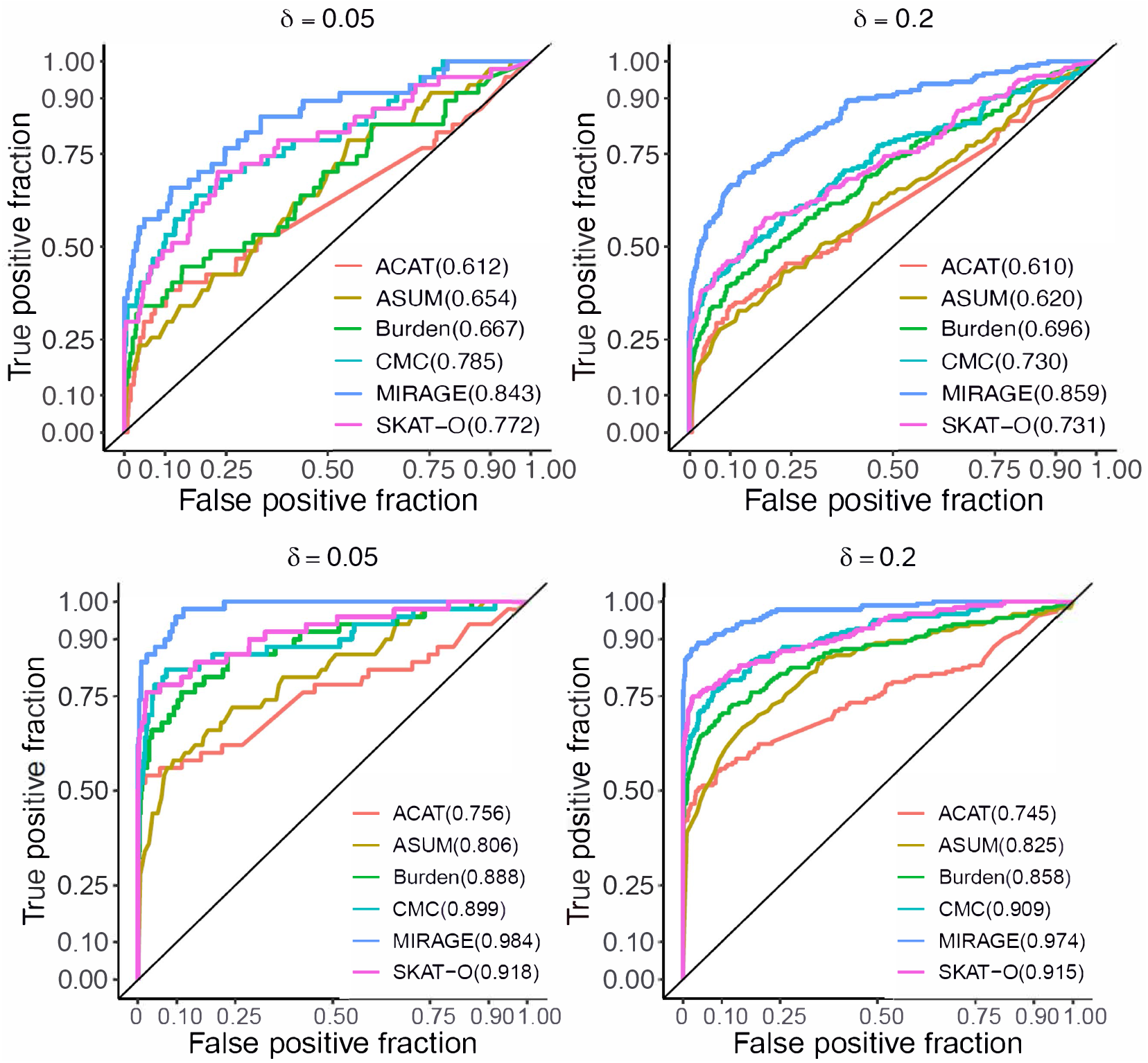
ROC curves of different methods for classifying risk genes. We simulate 1000 genes with varying proportion of risk genes, *δ* = 0.05, 0.2. AUC values are shown in the bracket. Solid black reference line is in the diagonal. The two figures at the top: results from using the same number of variants, i.e., 100 in a gene, and the bottom two figures: results from sampling random number of variants between 50 and 500 per gene.

**Figure S3.**
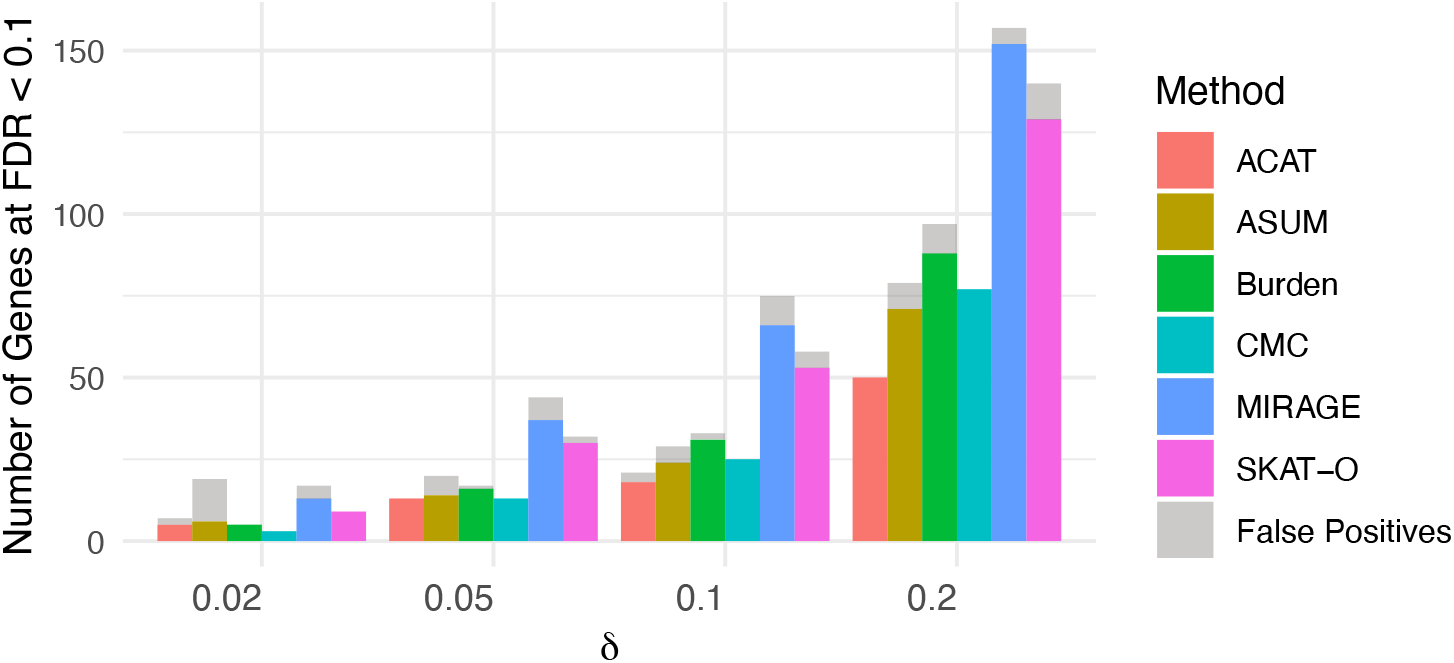
Number of true positive and false positive genes at FDR *<* 0.1 across different methods. The numbers of true positive and false positive genes for each *δ* and method are shown in the figure. Colored bars represent true positive genes, while grey bars indicate false positives

**Figure S4.**
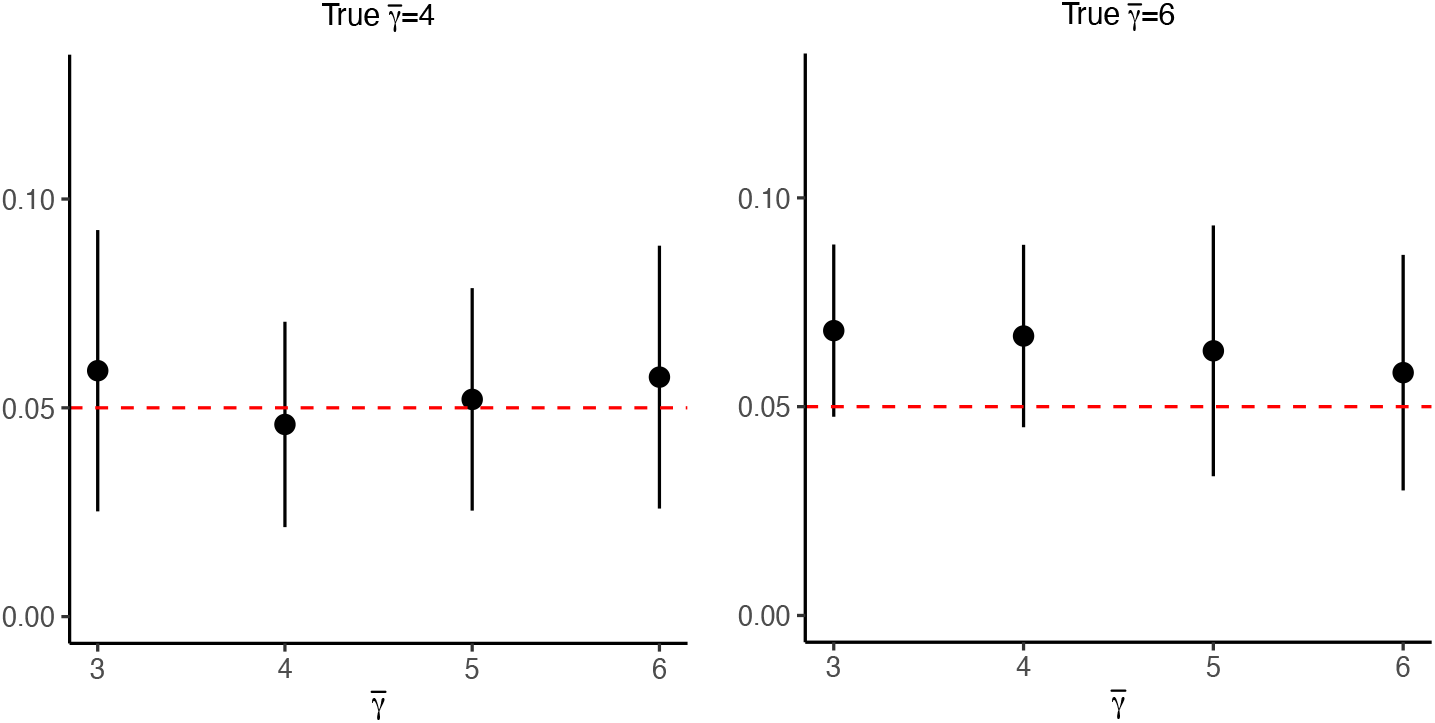
FDR calibration by MIRAGE. We simulated 20 datasets with true 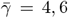, use mis-specified 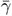 to calculate FDR. Red dashed line is the true FDR reference line.

**Figure S5.**
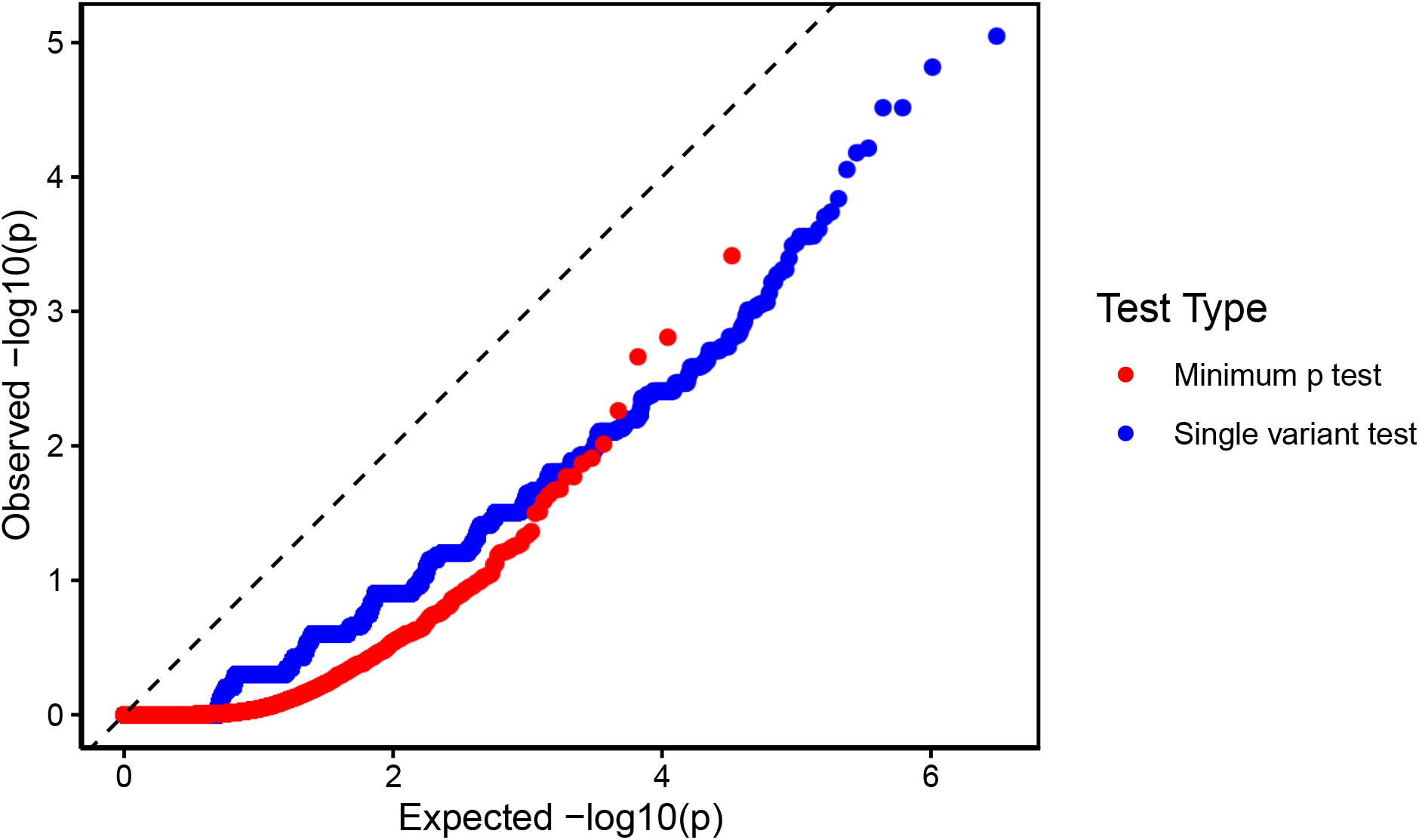
QQ plot of SNPs’ p values from the single variant test and genes’ p values from minimum-p test in the ASD dataset.

**Figure S6.**
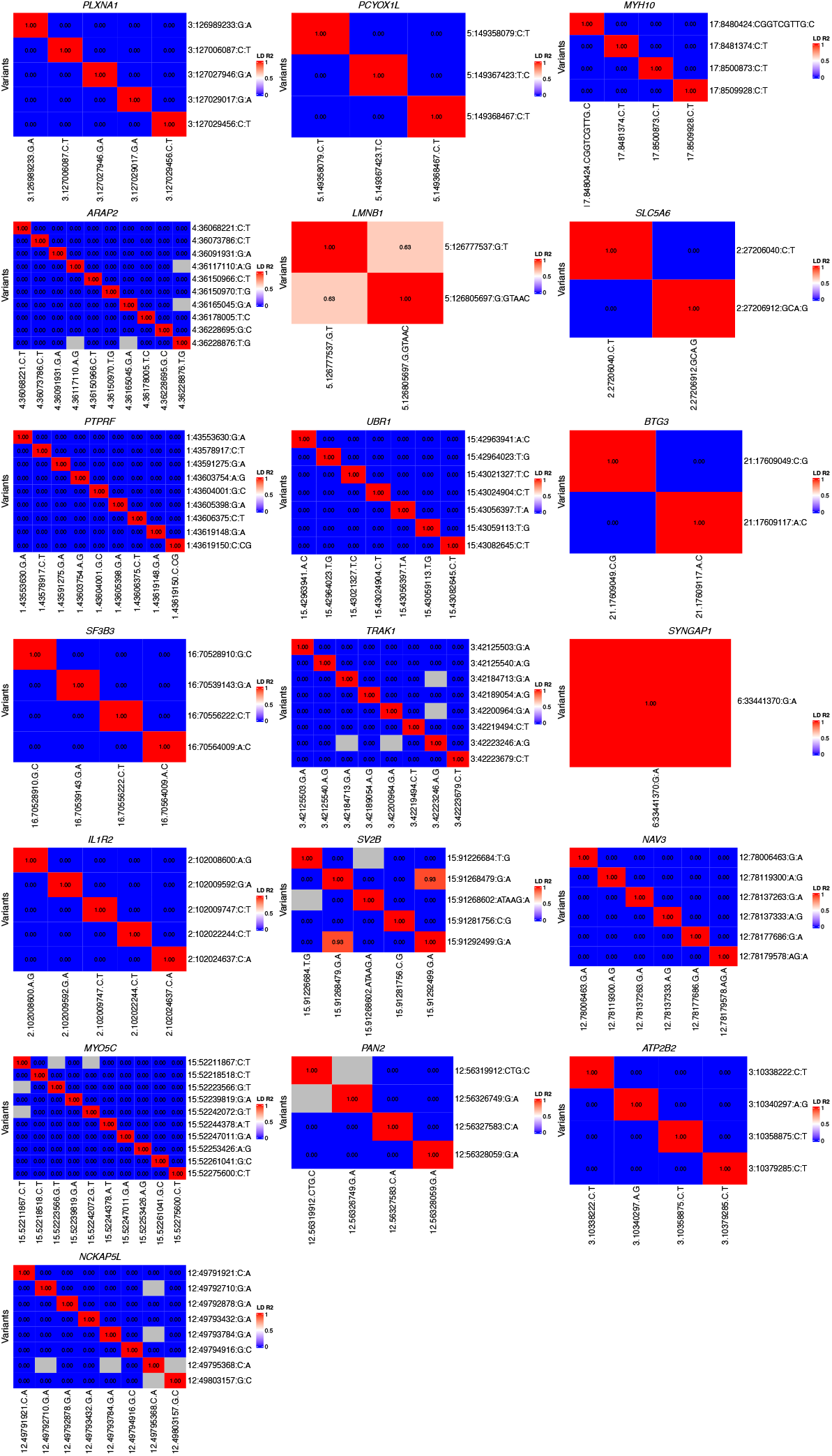
Heatmap showing LD (R^2^) of the supporting variants of genes having PP *>* 0.5. Supporting variants of a gene are defined as the variants whose logBF *>* 1. Gray elements show the variant pairs whose allele frequencies were too low to compute LD.

**Figure S7.**
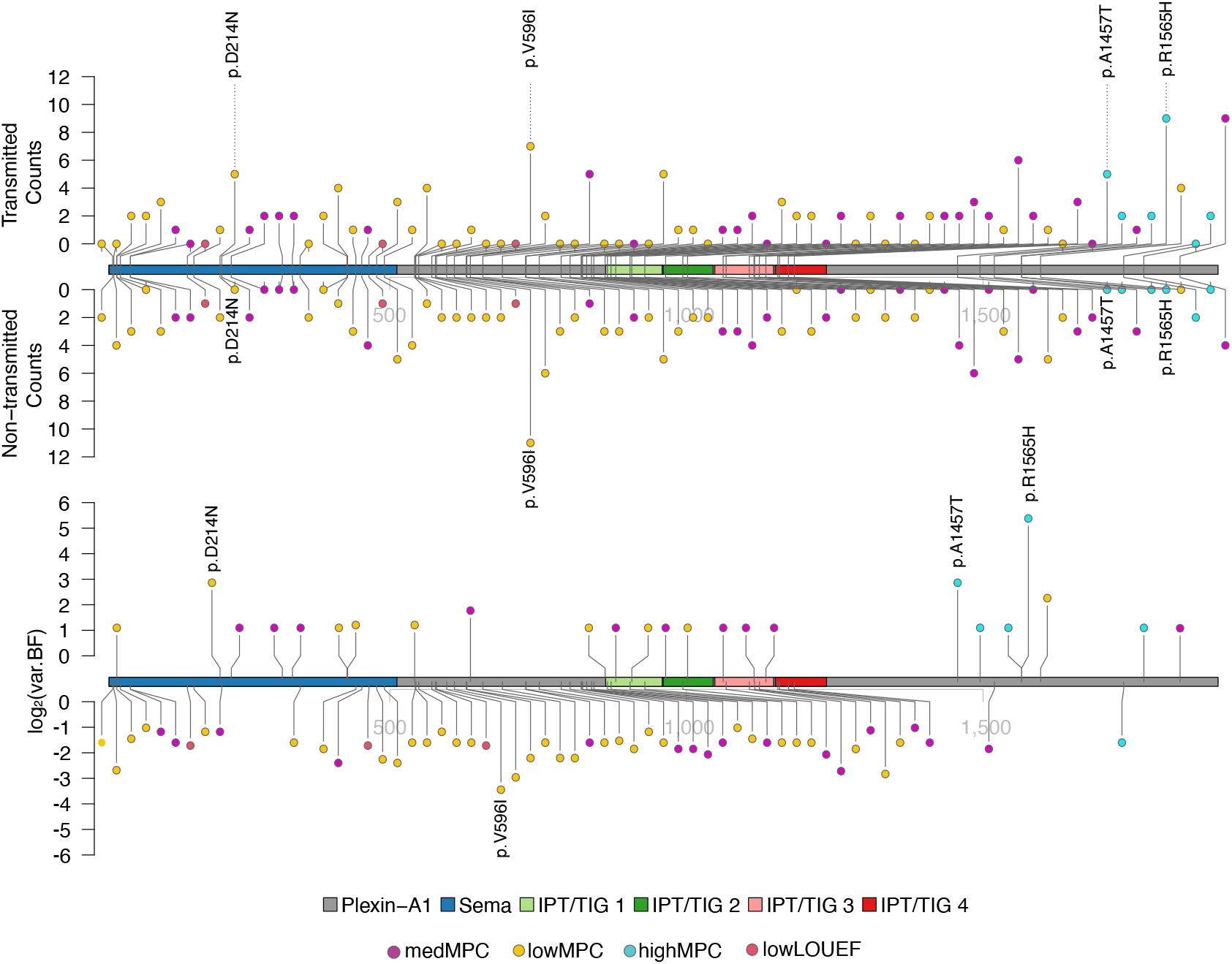
The lollipop plot of *PLXNA1*. Top: the transmission and non-transmission counts of each variant. Bottom: the evidence (log_2_BF) of each variant. Only informative variants, defined as BF *>* 2 or BF*<* 0.5, were plotted. The variants with BF *>* 5 and BF *<* 0.1 were labelled. LOEUF (Loss-Of-function Observed/Expected Upper bound Fraction). MPC (Missense badness, PolyPhen-2, and Constrain scores).

**Figure S8.**
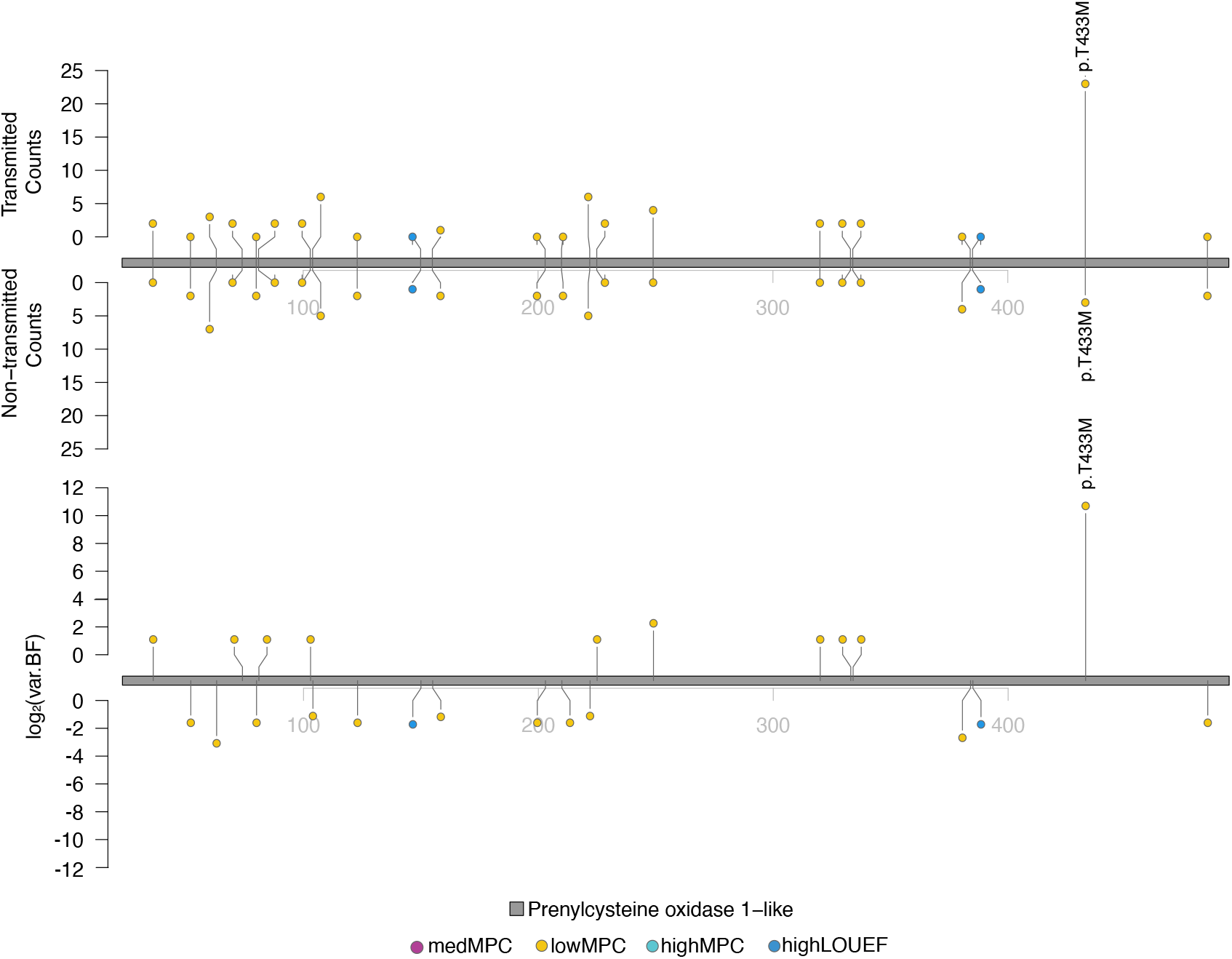
The lollipop plot of *PCYOX1L*. Top: the transmission and non-transmission counts of each variant. Bottom: the evidence (log_2_BF) of each variant. Only informative variants, defined as BF *>* 2 or BF*<* 0.5, were plotted. The variants with BF *>* 5 and BF *<* 0.1 were labelled. LOEUF (Loss-Of-function Observed/Expected Upper bound Fraction). MPC (Missense badness, PolyPhen-2, and Constrain scores).

**Figure S9.**
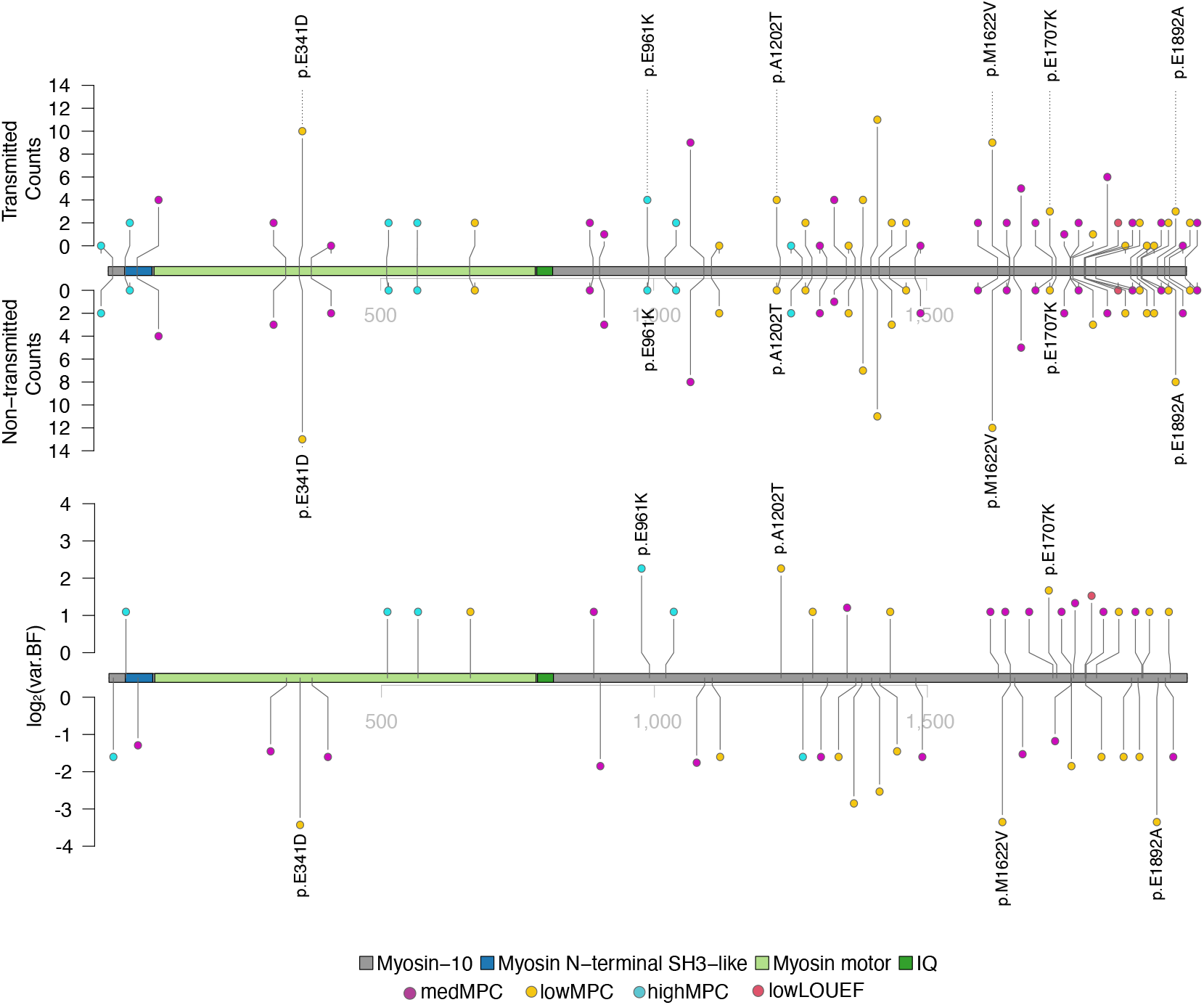
The lollipop plot of *MYH10*. Top: the transmission and non-transmission counts of each variant. Bottom: the evidence (log_2_BF) of each variant. Only informative variants, defined as BF *>* 2 or BF*<* 0.5, were plotted. The variants with BF *>* 3 and BF *<* 0.1 were labelled. LOEUF (Loss-Of-function Observed/Expected Upper bound Fraction). MPC (Missense badness, PolyPhen-2, and Constrain scores).

**Figure S10.**
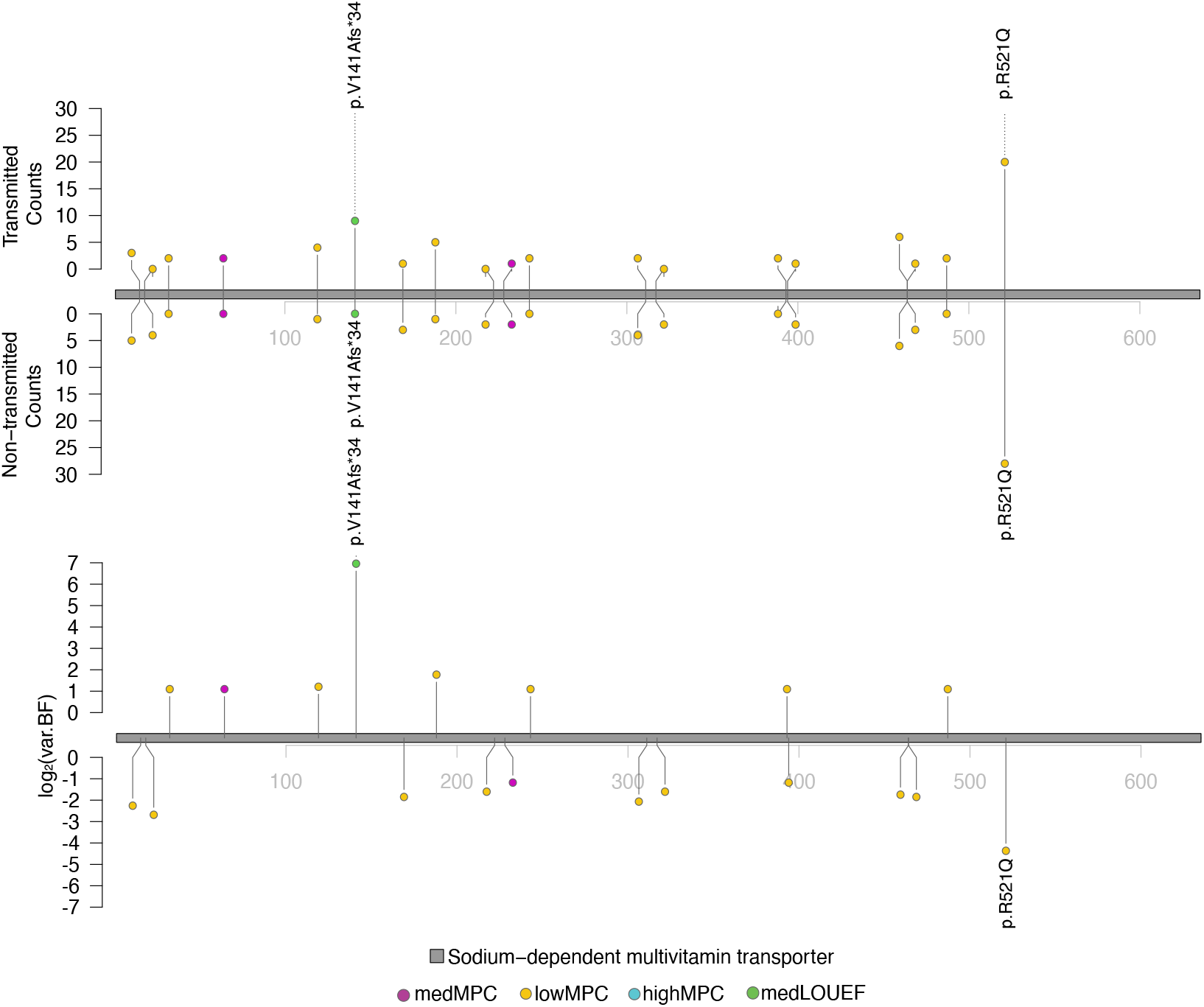
The lollipop plot of *SLC5A6*. Top: the transmission and non-transmission counts of each variant. Bottom: the evidence (log_2_BF) of each variant. Only informative variants, defined as BF *>* 2 or BF*<* 0.5, were plotted. The variants with BF *>* 3 and BF *<* 0.1 were labelled. LOEUF (Loss-Of-function Observed/Expected Upper bound Fraction). MPC (Missense badness, PolyPhen-2, and Constrain scores).

**Figure S11.**
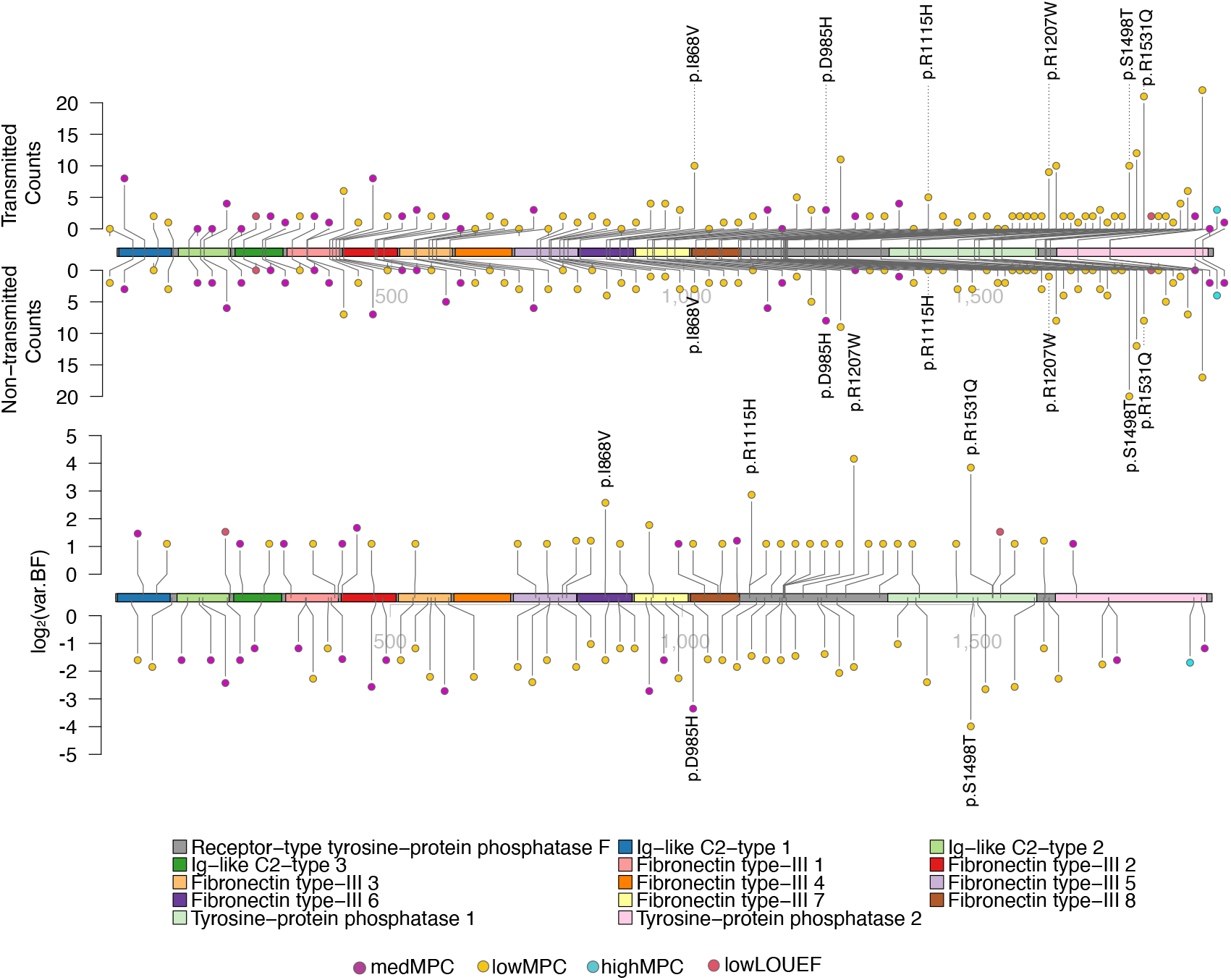
The lollipop plot of *PTPRF*. Top: the transmission and non-transmission counts of each variant. Bottom: the evidence (log_2_BF) of each variant. Only informative variants, defined as BF *>* 2 or BF*<* 0.5, were plotted. The variants with BF *>* 5 and BF *<* 0.1 were labelled. LOEUF (Loss-Of-function Observed/Expected Upper bound Fraction). MPC (Missense badness, PolyPhen-2, and Constrain scores).

**Figure S12.**
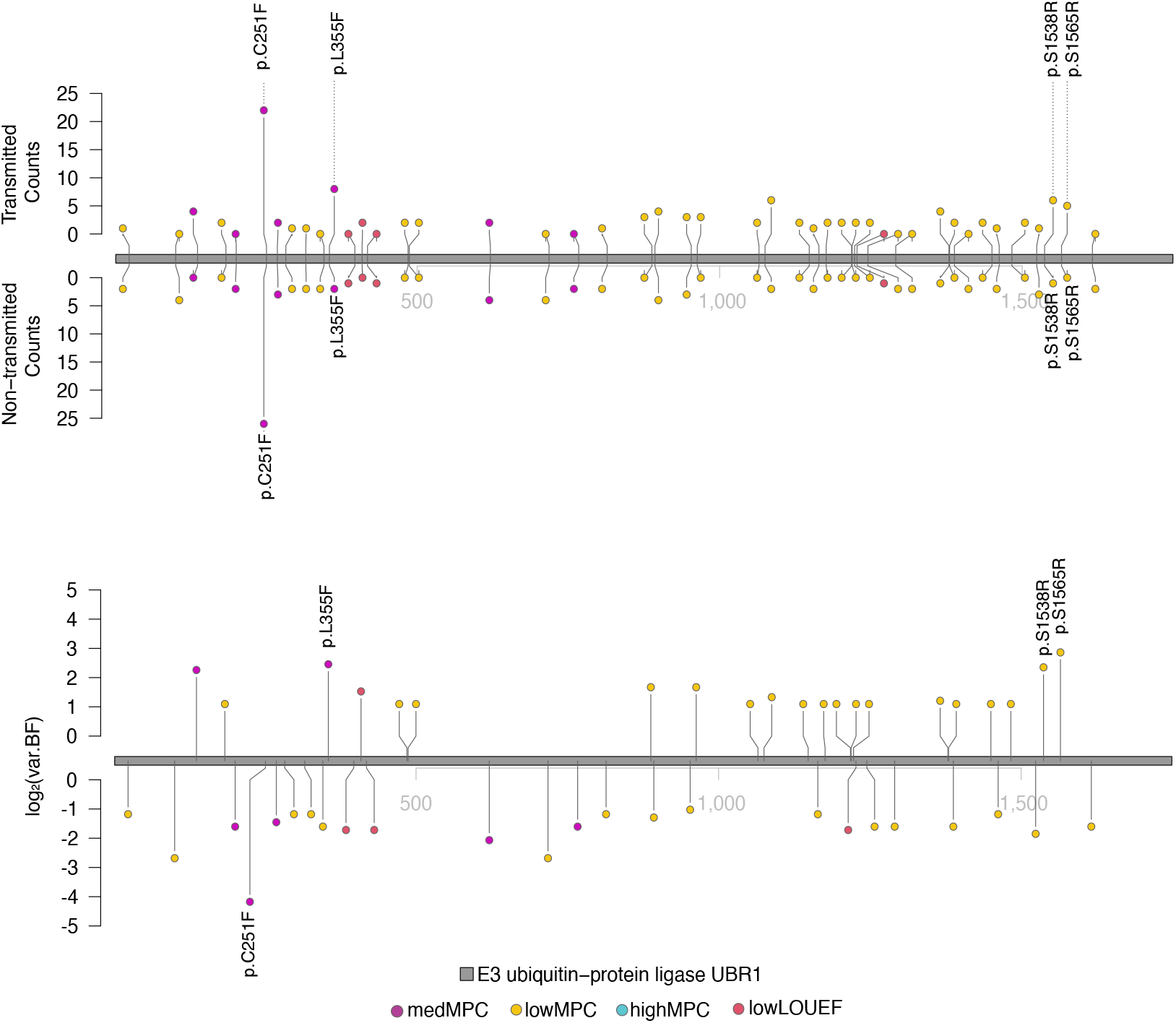
The lollipop plot of *UBR1*. Top: the transmission and non-transmission counts of each variant. Bottom: the evidence (log_2_BF) of each variant. Only informative variants, defined as BF *>* 2 or BF*<* 0.5, were plotted. The variants with BF *>* 5 and BF *<* 0.1 were labelled. LOEUF (Loss-Of-function Observed/Expected Upper bound Fraction). MPC (Missense badness, PolyPhen-2, and Constrain scores).

**Figure S13.**
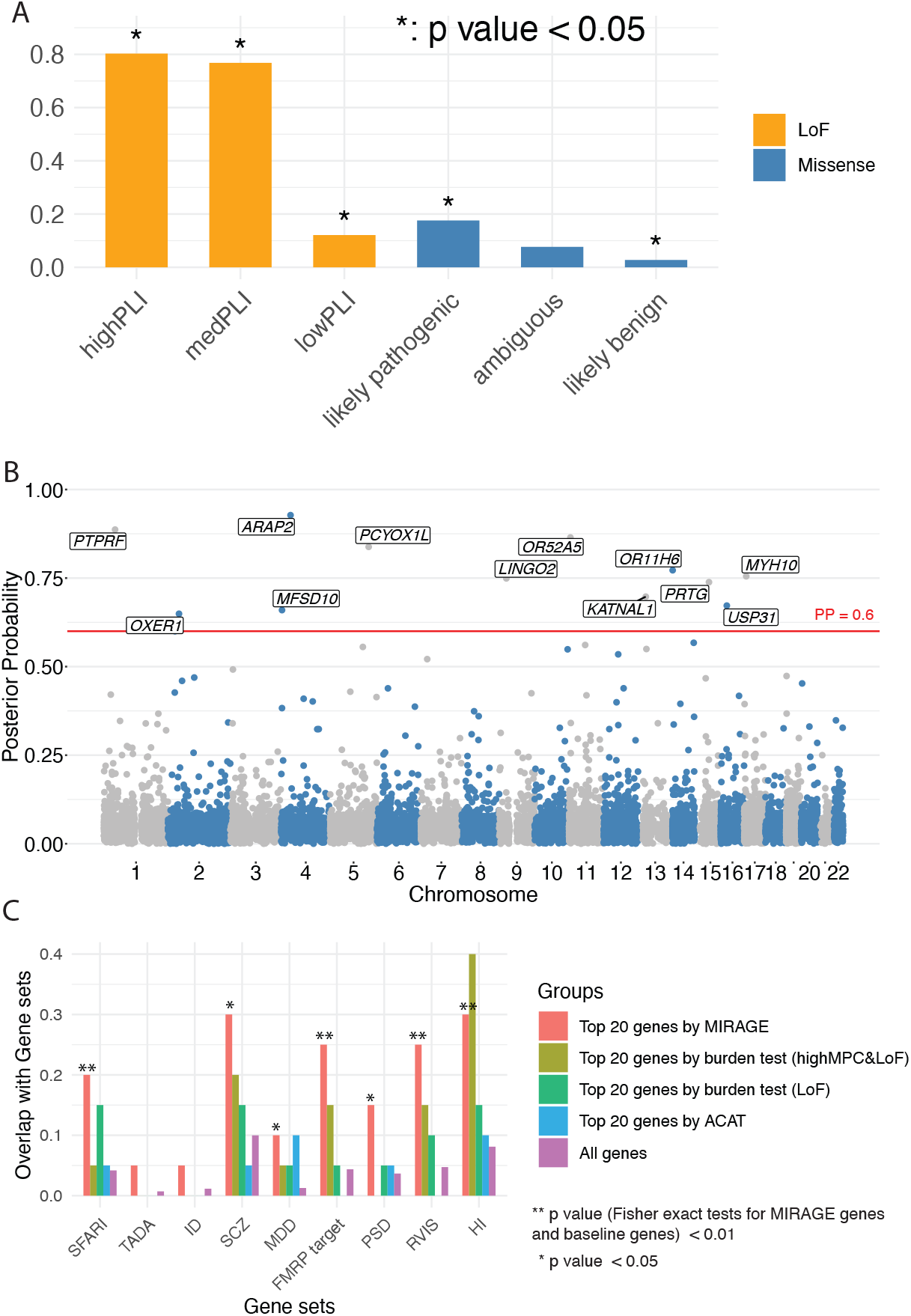
Results from using pLI as LoF annotation and AlphaMissense as missense variant annotation. **(A)** The proportion of risk variants in 6 variant categories. LoF categories: pLI (the probability of loss-of-function intolerance) ≥ 0.995 (high), 0.995 *>* pLI ≥ 0.5 (med), pLI *<* 0.5 (low). Missense categories: likely pathogenic, ambiguous, likely benign **(B)** Manhattan plot of posterior probabilities (PP) for all gene analyzed. Genes with PP *>* 0.6 are labeled. **(C)** Enrichment of ASD-related gene sets in the top 20 genes found by MIRAGE, burden tests and ACAT. Enrichment test was based on Fisher’s exact test. SFARI genes: Categories 1 and 2. TADA high confidence genes: FDR *<* 0.05. “All genes” refer to the entire set of genes analyzed, as a baseline.

**Figure S14.**
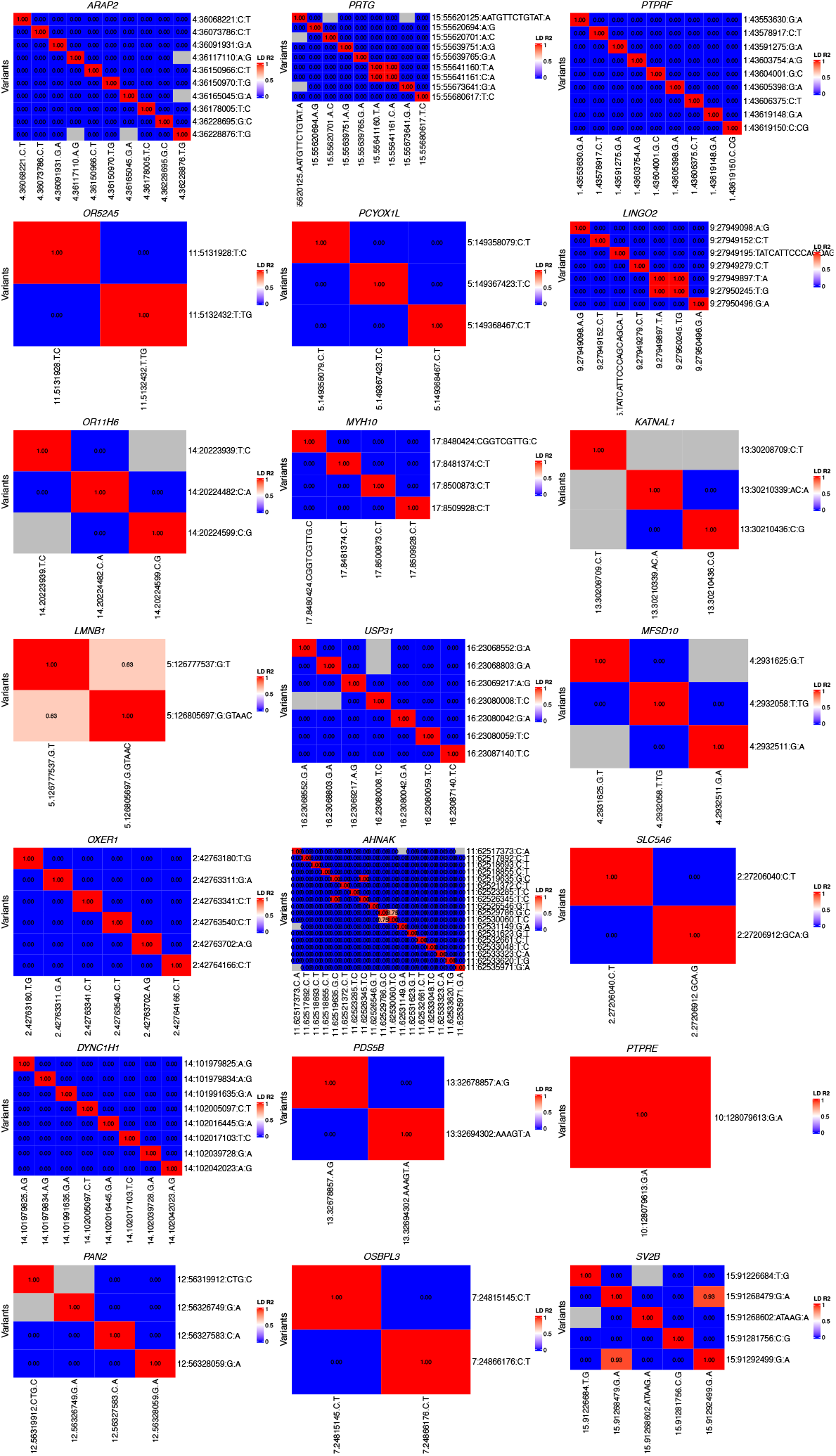
Heatmap showing LD (R^2^) of the supporting variants of genes having PP *>* 0.5. Variant annotations are pLI for LoF variants and AlphaMissense for missense variants. Supporting variants of a gene are defined as the variants whose logBF *>* 1. Gray elements show the variant pairs whose allele frequenci1e4s were too low to compute LD. pLI (the probability of loss-of-function intolerance).

**Figure S15.**
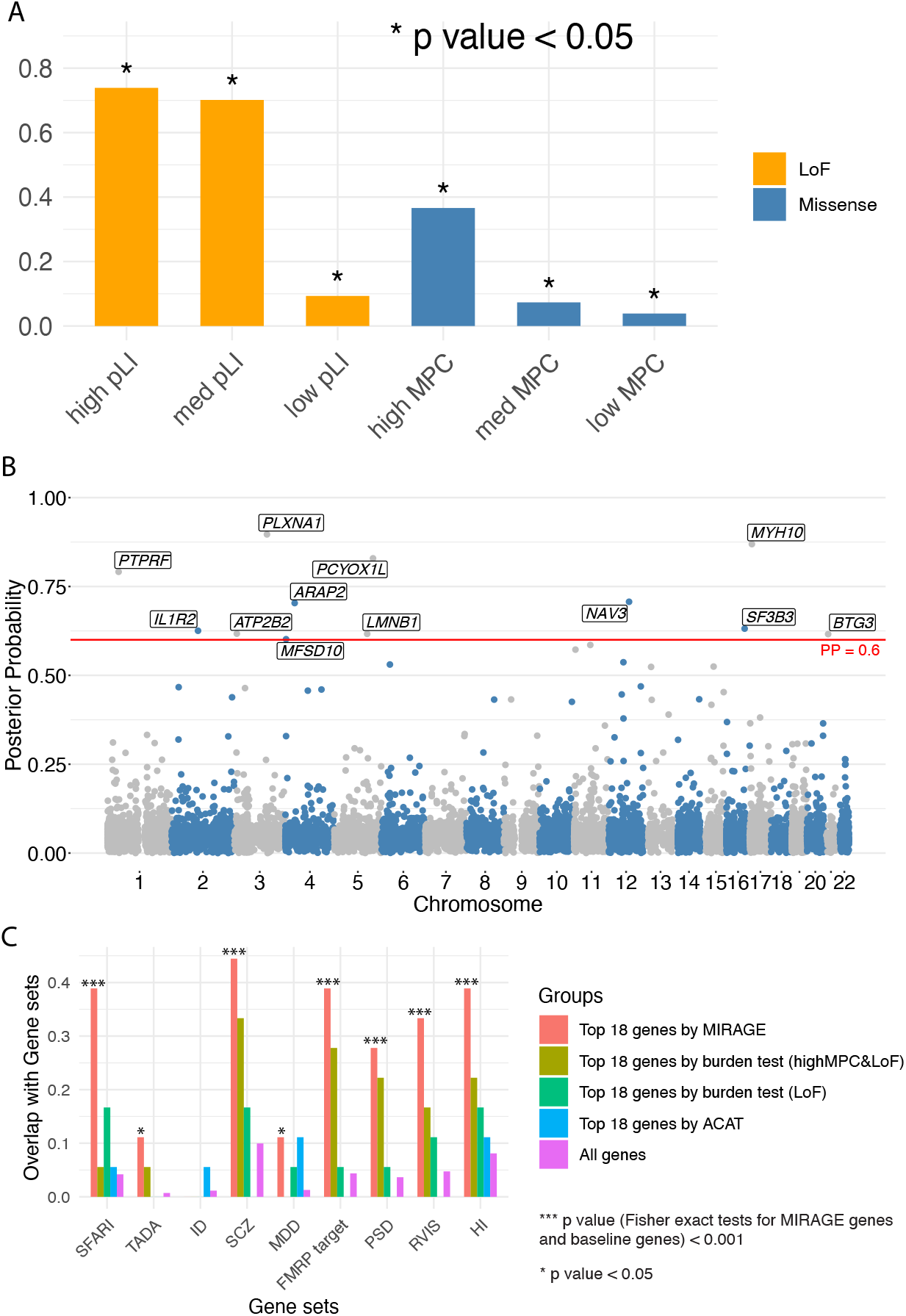
Results from using pLI as LoF annotation and MPC as missense variant annotation. **(A)** The proportion of risk variants in 6 variant categories. LoF categories: pLI ≥ 0.995 (high), 0.995 *>* pLI ≥ 0.5 (med), pLI *<* 0.5 (low). Missense categories: MPC≥ 2 (high), 2 *>* MPC ≥ 1 (med), MPC *<* 1 (low). **(B)** Manhattan plot of posterior probabilities (PP) for all gene analyzed. Genes with PP *>* 0.6 are labeled. **(C)** Enrichment of ASD-related gene sets in the top 18 genes found by MIRAGE, burden tests and ACAT. Enrichment test was based on Fisher’s exact test. SFARI genes: Categories 1 and 2. TADA high confidence genes: FDR *<* 0.05. “All genes” refer to the entire set of genes analyzed, as a baseline. pLI (the probability of loss-of-function intolerance). MPC (Missense badness **(D)** PolyPhen-2 **(E)** and Constrain scores).

**Figure S16.**
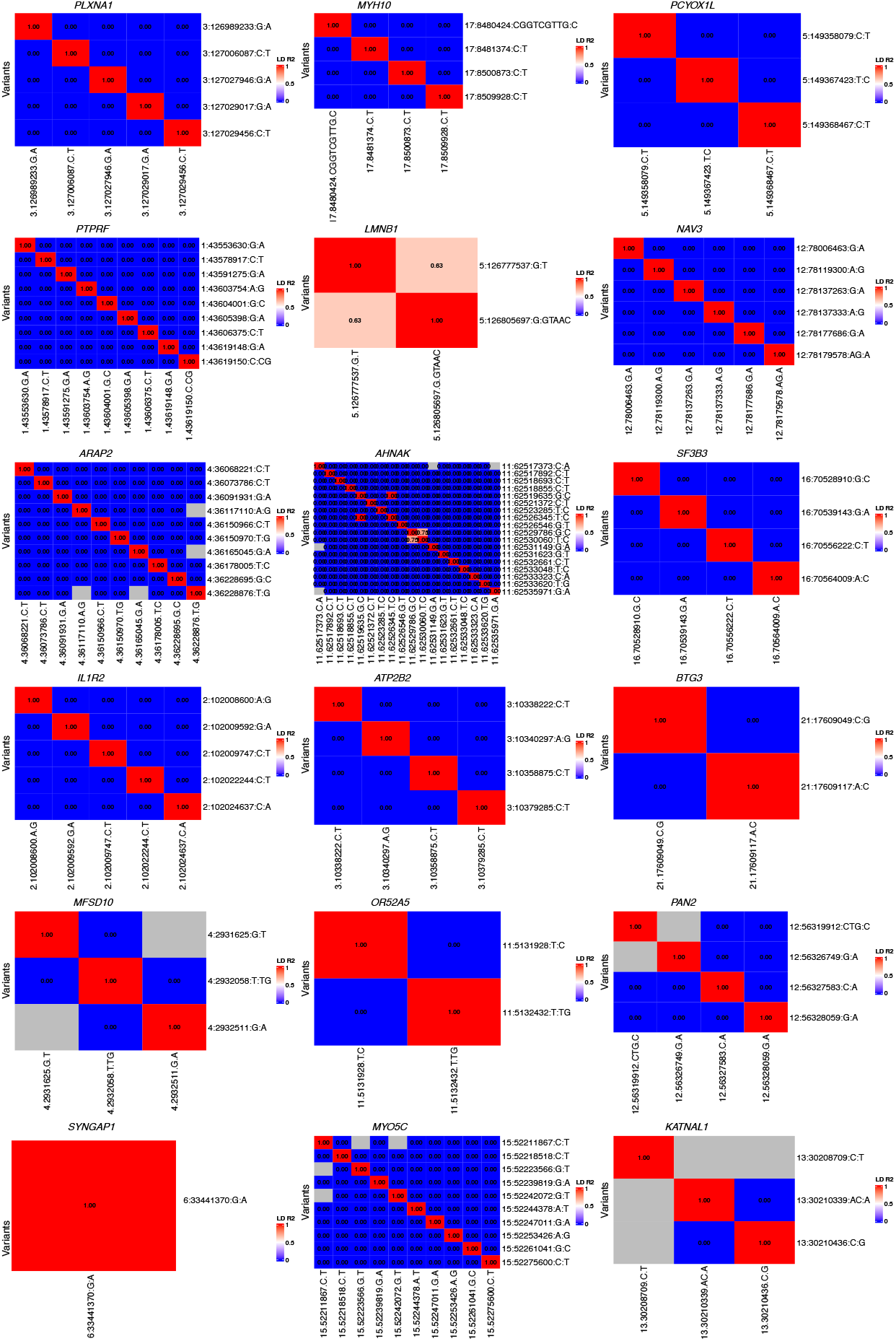
Heatmap showing LD (R^2^) of the supporting variants of genes having PP *>* 0.5. Variant annotations are pLI for LoF variants and MPC for missense variants. Supporting variants of a gene are defined as the variants whose logBF *>* 1. Gray elements show the variant pairs whose allele frequencies were too low to compute LD. pLI (the probability of loss-of-function intolerance).

## Supplemental Tables

**Table S1.**
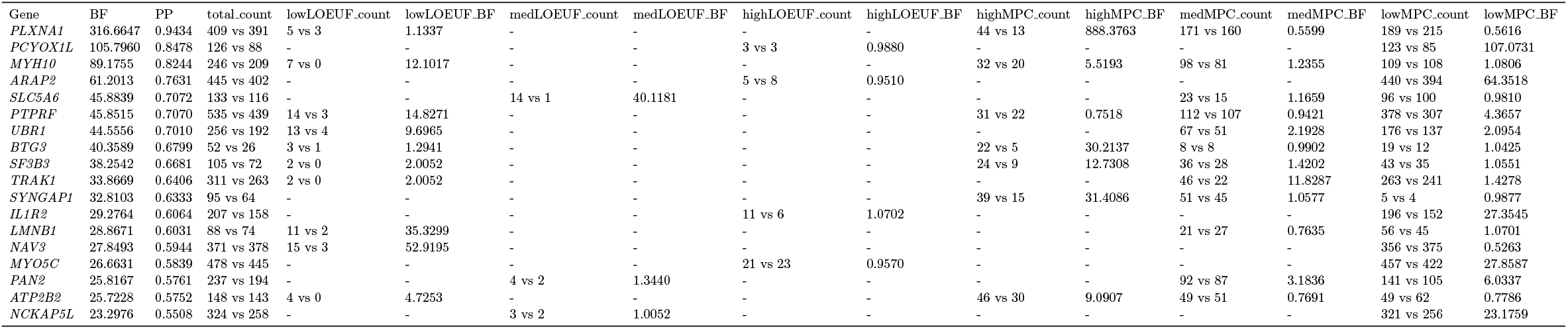
18 putative ASD risk genes identified by MIRAGE. BF (Bayesian Factor). PP (Posterior Probability). LOEUF (Loss-Of-function Observed/Expected Upper bound Fraction). MPC (Missense badness, PolyPhen-2, and Constrain scores).

| Gene | BF | PP | total_count | lowLOEUF_count | lowLOEUF_BF | medLOEUF_count | medLOEUF_BF | highLOEUF_count | highLOEUF_BF | highMPC_count | highMPC_BF | medMPC_count | medMPC_BF | lowMPC_count | lowMPC_BF |
| --- | --- | --- | --- | --- | --- | --- | --- | --- | --- | --- | --- | --- | --- | --- | --- |
| *PLXNA1* | 316.6647 | 0.9434 | 409 vs 391 | 5 vs 3 | 1.1337 | - | - | - | - | 44 vs 13 | 888.3763 | 171 vs 160 | 0.5599 | 189 vs 215 | 0.5616 |
| *PCYOX1L* | 105.7960 | 0.8478 | 126 vs 88 | - | - | - | - | 3 vs 3 | 0.9880 | - | - | - | - | 123 vs 85 | 107.0731 |
| *MYH10* | 89.1755 | 0.8244 | 246 vs 209 | 7 vs 0 | 12.1017 | - | - | - | - | 32 vs 20 | 5.5193 | 98 vs 81 | 1.2355 | 109 vs 108 | 1.0806 |
| *ARAP2* | 61.2013 | 0.7631 | 445 vs 402 | - | - | - | - | 5 vs 8 | 0.9510 | - | - | - | - | 440 vs 394 | 64.3518 |
| *SLC5A6* | 45.8839 | 0.7072 | 133 vs 116 | - | - | 14 vs 1 | 40.1181 | - | - | - | - | 23 vs 15 | 1.1659 | 96 vs 100 | 0.9810 |
| *PTPRF* | 45.8515 | 0.7070 | 535 vs 439 | 14 vs 3 | 14.8271 | - | - | - | - | 31 vs 22 | 0.7518 | 112 vs 107 | 0.9421 | 378 vs 307 | 4.3657 |
| *UBR1* | 44.5556 | 0.7010 | 256 vs 192 | 13 vs 4 | 9.6965 | - | - | - | - | - | - | 67 vs 51 | 2.1928 | 176 vs 137 | 2.0954 |
| *BTG3* | 40.3589 | 0.6799 | 52 vs 26 | 3 vs 1 | 1.2941 | - | - | - | - | 22 vs 5 | 30.2137 | 8 vs 8 | 0.9902 | 19 vs 12 | 1.0425 |
| *SF3B3* | 38.2542 | 0.6681 | 105 vs 72 | 2 vs 0 | 2.0052 | - | - | - | - | 24 vs 9 | 12.7308 | 36 vs 28 | 1.4202 | 43 vs 35 | 1.0551 |
| *TRAK1* | 33.8669 | 0.6406 | 311 vs 263 | 2 vs 0 | 2.0052 | - | - | - | - | - | - | 46 vs 22 | 11.8287 | 263 vs 241 | 1.4278 |
| *SYNGAP1* | 32.8103 | 0.6333 | 95 vs 64 | - | - | - | - | - | - | 39 vs 15 | 31.4086 | 51 vs 45 | 1.0577 | 5 vs 4 | 0.9877 |
| *IL1R2* | 29.2764 | 0.6064 | 207 vs 158 | - | - | - | - | 11 vs 6 | 1.0702 | - | - | - | - | 196 vs 152 | 27.3545 |
| *LMNB1* | 28.8671 | 0.6031 | 88 vs 74 | 11 vs 2 | 35.3299 | - | - | - | - | - | - | 21 vs 27 | 0.7635 | 56 vs 45 | 1.0701 |
| *NAV3* | 27.8493 | 0.5944 | 371 vs 378 | 15 vs 3 | 52.9195 | - | - | - | - | - | - | - | - | 356 vs 375 | 0.5263 |
| *MYO5C* | 26.6631 | 0.5839 | 478 vs 445 | - | - | - | - | 21 vs 23 | 0.9570 | - | - | - | - | 457 vs 422 | 27.8587 |
| *PAN2* | 25.8167 | 0.5761 | 237 vs 194 | - | - | 4 vs 2 | 1.3440 | - | - | - | - | 92 vs 87 | 3.1836 | 141 vs 105 | 6.0337 |
| *ATP2B2* | 25.7228 | 0.5752 | 148 vs 143 | 4 vs 0 | 4.7253 | - | - | - | - | 46 vs 30 | 9.0907 | 49 vs 51 | 0.7691 | 49 vs 62 | 0.7786 |
| *NCKAP5L* | 23.2976 | 0.5508 | 324 vs 258 | - | - | 3 vs 2 | 1.0052 | - | - | - | - | - | - | 321 vs 256 | 23.1759 |

**Table S2.**
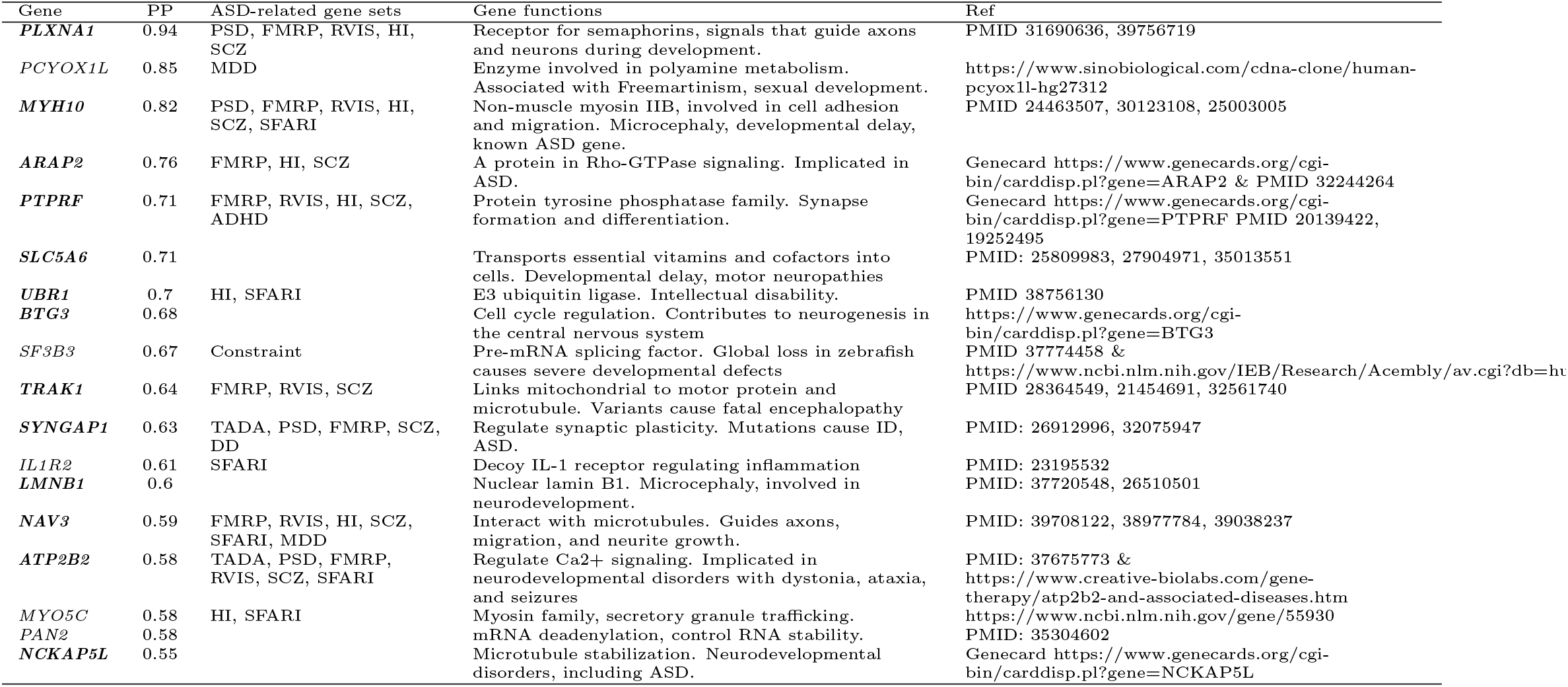
Genes identified by MIRAGE (posterior probability *>* 0.5, plausible genes are bolded) PP (Posterior Probability).

### EM Algorithm

We used Maximum Likelihood to estimate the parameters, denoted as *θ* = (*δ, η*), where *δ* is the proportion of risk genes and *η* (vector) the proportions of risk variants in each variant category. We note that the model has hyperparameters 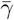 and *σ*. We assume the values are provided by the users, and found that in general, the results were relatively robust to the exact values (**Figures 2C–2D, Figure S4**). To estimate *θ*, we note that the model marginalizes latent variables **U** the risk gene status, and **Z** - the risk variant status. So we use Expectation-Maximization algorithm to estimate the parameters. We denote *θ*^(*t*)^ the parameter estimate at the *t*-th iteration.

- E step: we compute the *Q*(·) function

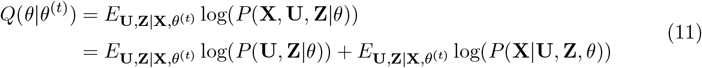

First we derive log(*P* (**U, Z**|*θ*)) and its expectations.

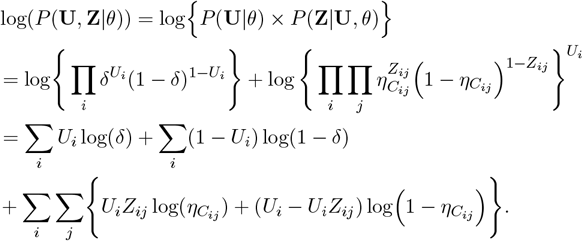

Taking expectation over **U, Z**|**X**, *θ*^(*t*)^:

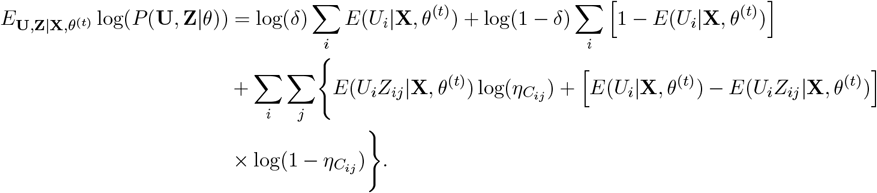

We will derive below the expectation terms *E*(*U*_*i*_|**X**, *θ*^(*t*)^) and *E*(*U*_*i*_*Z*_*ij*_|**X**, *θ*^(*t*)^).

Next we derive log(*P* (**X**|**U, Z**, *θ*)) and its expectation. Note that when gene *i* is a non-risk gene, i.e. *U*_*i*_ = 0, *P* (*X*_*i*_|*U*_*i*_ = 0, *θ*) is independent of *θ*, so

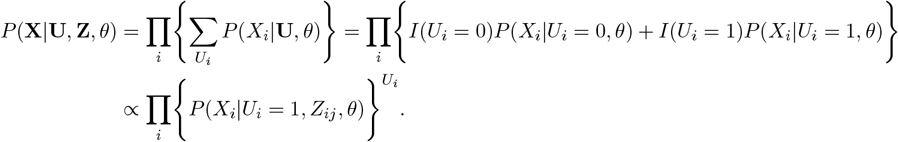

We expand *P* (*X*_*i*_|*U*_*i*_ = 1, *Z*_*ij*_, *θ*) into the variant level probabilities. Using the definition of *B*_*ij*_ in Equation 9, we have:

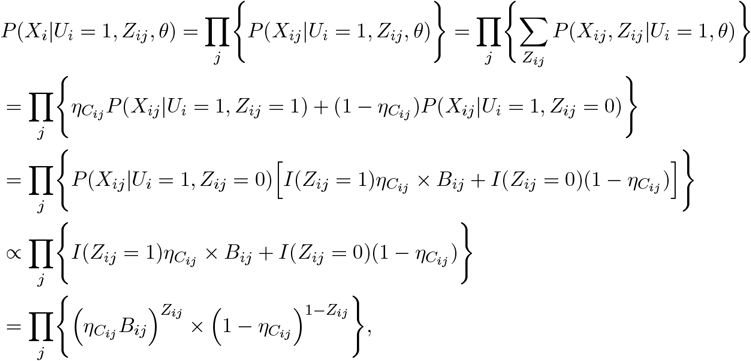

therefore,

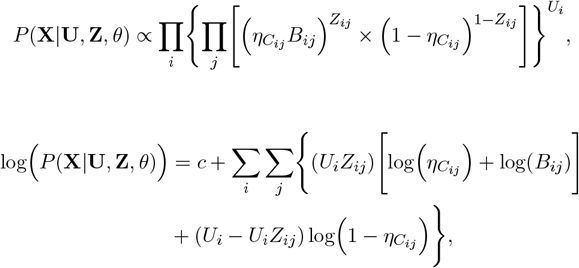

where *c* is a constant. Now we take the expectation over **U, Z**|**X**, *θ*^(*t*)^:

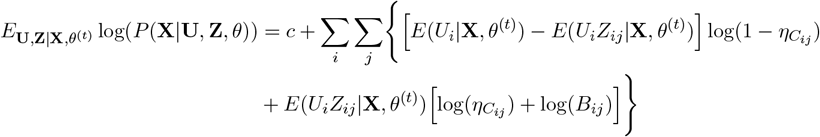

Putting them together, Equation 11 becomes:

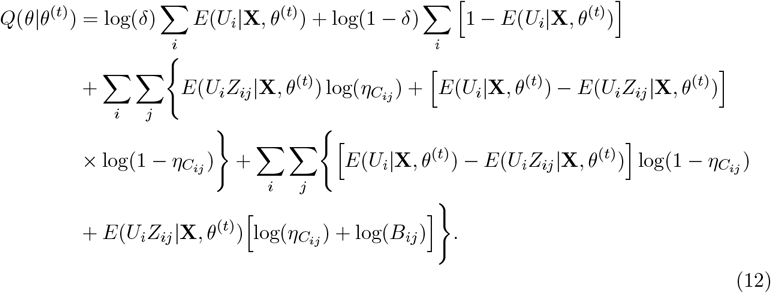

In the *Q* function, two expectations need to be calculated, *E*(*U*_*i*_|**X**, *θ*^(*t*)^), *E*(*U*_*i*_*Z*_*ij*_|**X**, *θ*^(*t*)^). First,

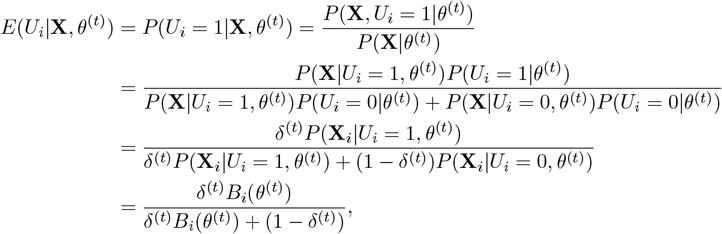

where *B*_*i*_ is the Bayes factor for gene *i*, as in Equation 8. Note that the BF of a gene depends on the parameters at the *t*-th iteration, namely *η*^(*t*)^. This term has a simple interpretation: the posterior probability of the gene *i* being a risk gene given the current parameters. The second expectation term also has a simple interpretation: the posterior probability that gene *i* is a risk gene and its variant *j* is a risk variant, given the current parameters. We denote *λ*_*ij*_ = *P* (*X*_*ij*_|*U*_*i*_ = 1, *Z*_*ij*_ = 0, *θ*), then *P* (*X*_*ij*_|*U*_*i*_ = 1, *Z*_*ij*_ = 1, *θ*) = *λ*_*ij*_ *B*_*ij*_,

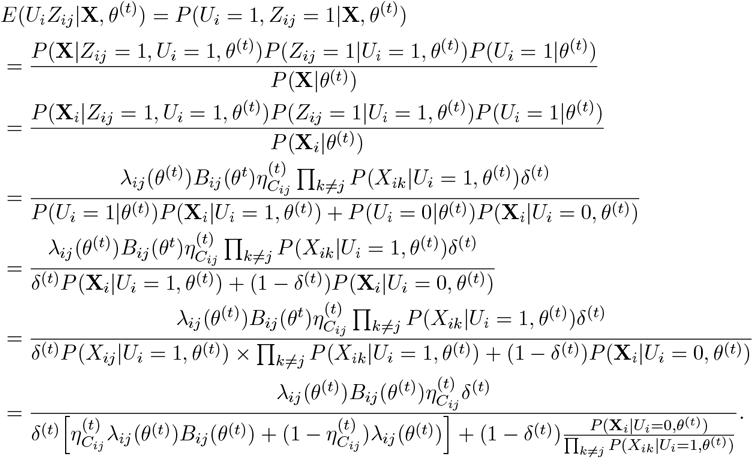

Note that

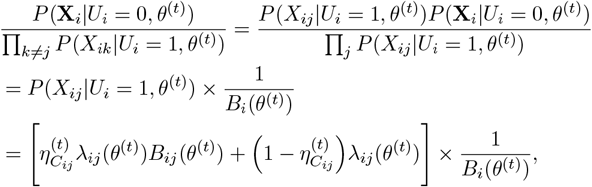

thus,

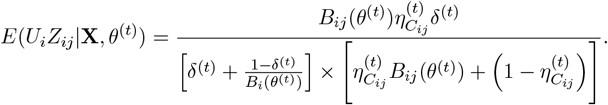

With these expectation terms, we completed our calculation of the *Q* function.

- M step: we update *θ* by *θ*^(*t*+1)^ that maximizes *Q*(*θ*|*θ*^(*t*)^) as,

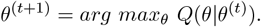

We first take derivative with respect to *δ*. Note that in Equation 12, *δ* only occurs in the first two terms, so we have:

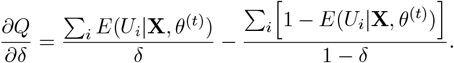

Setting this equations to 0 leads to:

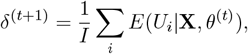

where *I* is the total number of genes.

Next we derive the update rule of *η*. Assume there are *C* annotation groups, then *η* = (*η*_1_, *η*_2_, …, *η*_*C*_) is a *C*-dimensional vector. The variant (*i, j*) in group *c* has the prior probability of being causal variant *η*_*c*_. We use *v*(*i, j*) to denote the group this variant belongs to. Taking the derivative of *Q* function with respect to *η*_*c*_, we have:

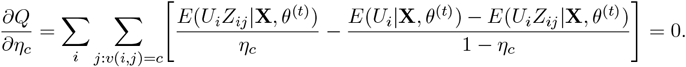

Solving this equation leads to the update rule:

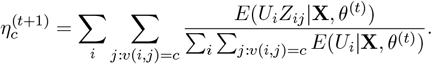

For parameters 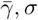, there are no closed solutions for 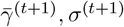. Instead, we fix 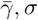 based on empirical evidence.

### Results using pLI as LoF annotation, AlphaMissense as missense annotation

In this section, when annotating variants, we used probability of loss-of-function intolerance (pLI scores) for LoF variants^38,39^, and AlphaMissense pathogenicity^42^ for missense variants. We included a total of six variant groups: pLI ≥ 0.995 (high pLI), 0.995 *>* pLI ≥ 0.5 (med pLI), pLI *<* 0.5 (low pLI), in addition, AlphaMissense classification: likely pathogenic, ambiguous, and likely benign.

To run MIRAGE, we used the same settings as the main text did. MIRAGE estimated the proportions of risk variants across the six variant categories (**Figure S13A**). The LoF variants in genes with high and medium pLI showed the highest proportions, with ∼ 80% of variants being risk variants. In the missense variant groups, high MPC had the largest risk variant proportion of ∼ 20%, and the last two categories showed much lower proportions. These estimates were generally in line with the expected deleteriousness of these variant annotations. With the estimated parameters, we calculated BFs and PP for every gene and performed Bayesian FDR control. At PP *>* 0.5, MIRAGE identified 20 putative ASD risk genes (**Figure S13B**). We verified that the vast majority of the variants supporting these genes have very low LD (**Figure S14**), and pruning of the few remaining variants in LD have small effects on the results. At PP *>* 0.7, MIRAGE identified nine putative ASD risk genes. These results thus supported higher sensitivity of MIRAGE in detecting risk genes than existing rare variant association methods.

We evaluated the candidate genes by assessing the enrichment of ASD-related gene sets. We selected the 20 genes at PP *>* 0.5, and for comparison, the same number of top genes by burden tests and ACAT. The ASD-related gene sets include known ASD genes curated by SFARI^44^ and from de novo mutation studies using TADA^36^; risk genes of Intellectual Disability (ID)^45^ and Schizophrenia (SCZ)^46^; relevant biological processes including Post-synaptic density (PSD)^46^ and FMRP target genes^46^; evolutionarily constrained genes from RVIS^47^, Haplo-insufficient (HI) genes^47^ and major depressive disorder^48^. We note that the SFARI and TADA genes were derived from independent datasets. While some samples in our dataset were included in earlier studies, the transmission data were not used previously. We found that the top MIRAGE genes were enriched in most ASD related gene sets. The top genes from other methods showed lower or no enrichment. These enrichment results thus strongly supported the likely roles of the candidate genes in ASD.

We next examined the function and plausibility of the top 9 genes using the more stringent cutoff, PP *>* 0.7). Among the nine genes, *PTPRF* and *ARAP2* have known or related functions in neurodevelopment and/or other neuropsychiatric disorders. For example, both of them have been implicated in SCZ. *MYH10* was a ASD risk gene according to SFARI (score 2, Strong candidate). It was supported by multiple *de novo* mutation studies^54–57^. *PRTG* has been reported as a candidate risk gene for ASD^94,95^. In mouse and chick models, it marks early neural progenitors and suppresses premature neuronal differentiation during neural development^96^. The soluble form of *LINGO2* promotes excitatory synapse formation and has been implicated in autism spectrum disorder (ASD) by multiple studies^97–99^. *KATNAL1* regulates neuronal development, migration, and behavior, with loss-of-function linked to brain abnormalities and ciliary defects in mice^100^.

### Results using pLI as LoF annotation, MPC as missense annotation

In this section, we used the same functional categories as in an earlier study^37^ to annotate variants, based on probability of loss-of-function intolerance (pLI scores) for LoF variants^38,39^, and MPC scores (Missense badness, PolyPhen-2, and Constrain scores) for missense variants^41^. We included a total of six variant groups: pLI ≥ 0.995 (high pLI), 0.995 *>* pLI ≥ 0.5 (med pLI), pLI *<* 0.5 (low pLI), MPC≥ 2 (high MPC), 2 *>* MPC ≥ 1 (med MPC) and MPC *<* 1 (low MPC).

To run MIRAGE, we used the same settings as the main text did. MIRAGE estimated the proportions of risk variants across the six variant categories (**Figure S15A**). The LoF variants in genes with high and medium pLI showed the highest proportions, with ∼ 70% of variants being risk variants. In the missense variant groups, high MPC had the largest risk variant proportion of ∼ 37%, and the last two categories showed much lower proportions. These estimates were generally in line with the expected deleteriousness of these variant annotations.

We evaluated the candidate genes by assessing the enrichment of ASD-related gene sets (**Figure S15C**). We selected the 18 genes at PP *>* 0.5 (**Figure S15B**), and for comparison, the same number of top genes by burden tests and ACAT. The ASD-related gene sets include known ASD genes curated by SFARI^44^ and from de novo mutation studies using TADA^36^; risk genes of Intellectual Disability (ID)^45^ and Schizophrenia (SCZ)^46^; relevant biological processes including Post-synaptic density (PSD)^46^ and FMRP target genes^46^; evolutionarily constrained genes from RVIS^47^, Haplo-insufficient (HI) genes^47^ and major depressive disorder^48^. We note that the SFARI and TADA genes were derived from independent datasets. While some samples in our dataset were included in earlier studies, the transmission data were not used previously. We found that the top MIRAGE genes were enriched in most ASD related gene sets. The top genes from other methods showed lower or no enrichment. These enrichment results thus strongly supported the likely roles of the candidate genes in ASD.

We next examined the function and plausibility of the top 6 genes using the more stringent cutoff, PP *>* 0.7). Among the six genes, *PLXNA1* and *NAV3* were found by in earlier ASD studies^34,36^. *MYH10* was a ASD risk gene according to SFARI (score 2, Strong candidate). It was supported by multiple *de novo* mutation studies *PTPRF* and *ARAP2* have known or related functions in neurodevelopment and/or other neuropsychiatric disorders. For example, both of them have been implicated in SCZ.

